# A glycine zipper motif governs translocation of type VI secretion toxic effectors across the cytoplasmic membrane of target cells

**DOI:** 10.1101/2022.07.12.499750

**Authors:** Jemal Ali, Manda Yu, Li-Kang Sung, Yee-Wai Cheung, Erh-Min Lai

## Abstract

Type VI secretion systems (T6SSs) can deliver diverse toxic effectors into eukaryotic and bacterial cells. Although much is known about the regulation and assembly of T6SS, the translocation mechanism of effectors into the periplasm and/or cytoplasm of target cells remains elusive. Here we use the *Agrobacterium tumefaciens* DNase effector Tde1 to unravel the mechanism of translocation from attacker to prey. We demonstrate that Tde1 binds to its adaptor Tap1 through the N-terminus, which harbours continuous copies of GxxxG motifs resembling the glycine zipper structure found in proteins involved in the membrane channel formation. Amino acid substitutions on G^39^xxxG^43^ motif does not affect Tde1-Tap1 interaction and secretion but abolish its membrane permeability and translocation of its fluorescent fusion protein into prey cells. The data suggest that G^39^xxxG^43^ governs the delivery of Tde1 into target cells by permeabilizing the cytoplasmic membrane. Considering the widespread presence of GxxxG motifs in bacterial effectors and pore-forming toxins, we propose that glycine zipper mediated permeabilization is a conserved mechanism used by bacterial effectors for translocation across target cell membranes.

## INTRODUCTION

In a complex microbial community, bacteria have evolved versatile secretion systems for export or import of substrates across their membranes in response to different environmental cues. Each specialized protein secretion systems (type I to X secretion system [TISS to TXSS])(reviewed in <u>Christie, 2019</u>; <u>Costa et al., 2015</u>; <u>Palmer et al., 2020</u>) can recognize specific substrates for secretion and translocation across one or multiple membranes. The type VI secretion system (T6SS) is a molecular weapon deployed by many Proteobacteria for pathogenesis, antagonism, or nutrient acquisition (<u>Coulthurst, 2019</u>). The T6SS effectors discovered so far exert functions in antibacterial, anti-eukaryotic, and metal acquisition (<u>Hachani et al., 2016</u>; <u>Jurenas and Journet, 2021</u>; <u>Lien and Lai,</u> <u>2017</u>; <u>Russell et al., 2014</u>). The most established T6SS effectors are bacterial toxins, in which bacteria also produce cognate immunity proteins to prevent self-intoxication and toxicity in the sibling cells.

T6SS is a multiprotein complex, composed of at least 13 conserved core proteins resembling a phage tail structure, that extends from the cytoplasm to the outer membrane of the attacker cell (<u>Cherrak et al., 2019</u>; <u>Wang et al., 2019</u>). The T6SS machine consists of the Tss(J)LM membrane complex (MC), TssEFGK base plate (BP), TssBC contractile sheath, and Hcp-VgrG-PAAR puncturing device. The MC interacts with the BP (<u>Cherrak et al., 2018</u>; <u>Durand et al., 2015</u>), which serves as a docking site of VgrG-PAAR effector complex to initiate polymerization of the tail (<u>Zoued et al., 2016</u>). The tail is composed of the Hcp inner tube and TssBC outer sheath, whose biogenesis is regulated by TssA cap protein, and when triggered, the sheath contracts and ejects out the effector decorated puncturing device into extracellular milieu or target cells (<u>Ali and Lai, 2022</u>; <u>Basler et al., 2012</u>; <u>Vettiger and</u> <u>Basler, 2016</u>).

The T6SS has multiple strategies for delivering diverse effectors. On the basis of the known effectors and their transport mechanisms, effectors can be classified as “specialized” or “cargo” effectors (<u>Cherrak</u> *<u>et al.</u>*<u>, 2019</u>; <u>Cianfanelli et al., 2016</u>). Specialized effectors are fused to either of the C-termini of three core structural proteins (Hcp, VgrG, or PAAR) while cargo effectors interact directly or require a specific chaperone/adaptor to be loaded into the lumen of the Hcp tube or onto the VgrG spike prior to secretion. Though diverse T6SS antibacterial effectors that act in the cytoplasm, membrane, or periplasm of the target cells have been reported (<u>Jurenas and Journet, 2021</u>; <u>Lien and Lai, 2017</u>; <u>Russell</u> *<u>et al.</u>*<u>, 2014</u>), their mechanisms to breach outer and inner membranes for targeting cytoplasm of their targets still yet to be clarified.

A glycine zipper structure consisting of repetitive GxxxG motifs is commonly found in membrane-associated proteins (<u>Kim et al., 2005</u>) and bacterial toxins (<u>Fonte et al., 2011</u>; <u>Kim</u> <u>et al., 2004</u>). Glycine zipper motifs are known be involved in toxicity of some bacterial effectors for membrane channel formation. For example, the transmembrane domain (TMD) of a vacuolating toxin, VacA of *Helicobacter pylori* encodes three GxxxG motifs forming helix-helix packing interactions (<u>Kim</u> *<u>et al.</u>*<u>, 2004</u>), which are required for the vacuolation and membrane channelling contributing to VacA toxicity (<u>McClain et al., 2003</u>). Type I secretion effectors CdzC and CdzD of *Caulobacter crescentus* and T6SS effector Tse4 of *Pseudomonas aeruginosa,* also possess glycine zipper motifs involved in antibacterial activity (<u>Garcia-Bayona et al., 2017</u>; <u>LaCourse et al., 2018</u>). Expression of Tse4 disrupted the proton motive force of the inner membrane while CdzC and CdzD form surface aggregation for contact-dependent killing of target cells. However, how glycine zipper motifs of Tse4 and CdzCD involved in toxicity remain unknown.

A T6SS-encoding locus is highly conserved in the genome of plant pathogenic bacterium *Agrobacterium tumefaciens* and the apparatus functions as an antibacterial weapon (<u>Chou</u> <u>et al., 2022</u>; <u>Ma et al., 2014</u>; <u>Wu et al., 2021</u>; <u>Yu et al., 2020</u>). We previously revealed that *A. tumefaciens* strain C58 deploys two <u>T</u>ype VI <u>D</u>Nase <u>e</u>ffectors (Tde1 and Tde2) as the major antibacterial weapons, in which the cognate immunity proteins (namely Tdi1 and Tdi2) prevent autointoxication (<u>Ma</u> *<u>et al.</u>*<u>, 2014</u>). Both Tde1 and Tde2 harbour a novel C-terminal Ntox15 toxin domain (<u>Zhang et al., 2012</u>) containing an HxxD catalytic motif required for its DNase activity (<u>Ma et al., 2014</u>). Tde1 requires its cognate chaperone/adaptor Tap1 for loading onto VgrG1 for secretion (<u>Bondage et al., 2016</u>).

By obtaining the uncoupling Tde1 variants that remain capable for binding to Tap1 for export but deficient in membrane permeability, translocation, and interbacterial competition, we reveal the secretion and translocation mechanism of Tde1 from the attacker cell to the target cell. We show that the N-terminal region of Tde1 harbouring repetitive glycine zipper motifs is sufficient for interacting with Tap1 for secretion. Once secreted, a conserved glycine zipper motif is necessary for translocation across target cell membranes. The findings demonstrate a new role of glycine zipper motif(s) in effector delivery into target cells.

## RESULTS

### Tde1 can cause DNase-independent growth inhibition in *Escherichia coli*

Our previous study showed that overexpression of Tde1 in *A. tumefaciens* C58 caused growth inhibition, and the immunity protein Tdi1 only partially protected against this cytotoxicity (<u>Ma</u> *<u>et al.</u>*<u>, 2014</u>). We hypothesized that Tde1 has domains apart from the DNase domain that contribute to its toxicity. In addition to the C-terminal Ntox15 DNase domain (amino acid 99-247) (<u>Ma</u> *<u>et al.</u>*<u>, 2014</u>), Tde1 has a predicted transmembrane domain (TMD, 22-42) (Fig.1A). Thus, three fragments of Tde1, that being the N-terminal, N-Tde1(1-97) and two C-terminal regions, C1-Tde1(49-278), and C2-Tde1(98-278) were tested for toxicity. To avoid confounding effects by the DNase activity, substitutions of catalytic residues (H190A, D193A) were introduced in the C1-Tde1 and the full length wide-type Tde1 to become C1-Tde1(M) and Tde1(M) respectively (Fig.1A). Ectopic expression in *E. coli* (DH10B) under an IPTG inducible promoter showed that N-Tde1 was sufficient to inhibit growth (Fig.1B). Tde1(M), but not the C1-Tde1(M), is growth inhibitory. Although C2-Tde1(WT) retains the wild type DNase catalytic residues, it was not able to inhibit growth, suggesting the N-terminus is required for the DNase activity. Both C1-Tde1(M) and C2-Tde1(WT) are expressed at levels similar or higher than N-Tde1 or Tde1(M), indicating that their loss of growth inhibition is not due to non-expression of the proteins (Fig. S1A). Evidence suggests the N-terminal region of Tde1 is sufficient to confer toxicity under conditions tested and that the C-terminal DNase domain requires entire or part of the N-terminus for it to cause toxicity.

**Figure 1.**
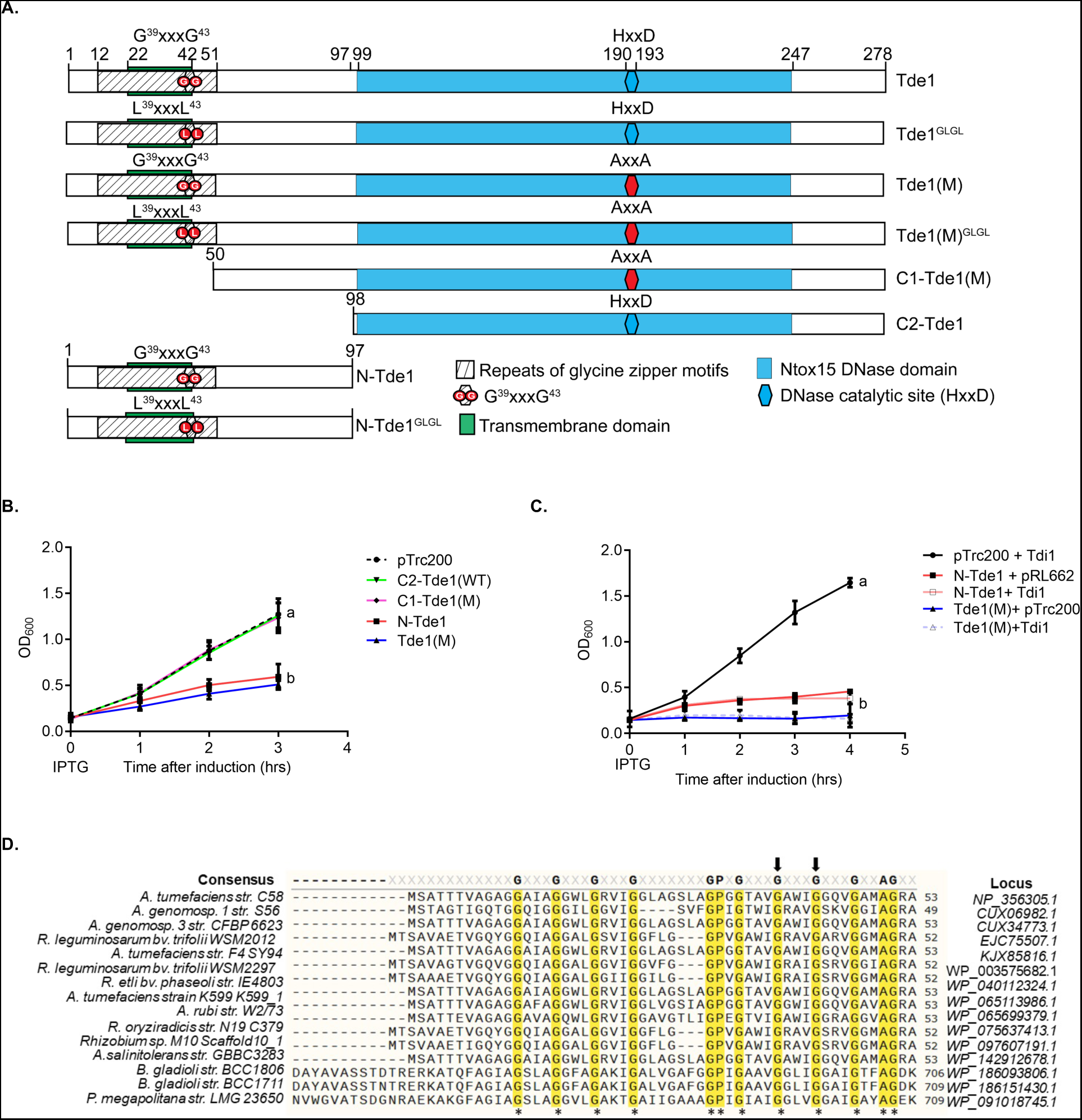
Schematic domain organization, sequence alignment, growth inhibition assay of Tde1. (A) Schematic domain organization of Tde1 protein and its variants. The N-terminal repeated glycine zipper motifs (12-51) overlapping a predicted transmembrane domain (22-42) and Ntox15 DNase domain (99-247) are indicated. Tde1 and its variants with truncation or amino acid substitutions were illustrated. (B) Growth inhibition assay of *E. coli* DH10B cells harbouring pTrc200 vector or each of its derivatives expressing Tde1 variants with IPTG induction. (C) Growth inhibition assay of *E. coli* DH10B cells co-expressing the Tde1 variants expressed from pTrc200 plasmid and Tdi1 immunity gene expressed from pRL662 plasmid. Growth curve was determined at OD_600._ Graphs of panels B and C show mean ± SD of three independent experiments. Different letters indicate statistically different groups of strains (p value <0.01). (D) Multiple sequence alignment of N-Tde1 homologues were presented with highly conserved amino acid residues highlighted in yellow. The bacterial species, strain name, and locus number of Tde1 orthologs (*Agrobacterium/Rhizobium*) or tape measure proteins (*Paraburkholderia/Burkholderia*) are indicated on the left and right of aligned sequences. Two conserved glycine residues (G^39^, G^43^) subjected for mutagenesis were indicated by the arrows above the sequences.

To test whether Tdi1, the immunity protein for the DNase toxicity of Tde1 (<u>Ma</u> *<u>et al.</u>*<u>, 2014</u>), can also neutralize the N-Tde1 toxicity, the Tde1 variants were co-expressed with the Tdi1. The result show that Tdi1 could not rescue the growth inhibition caused by the N-Tde1 and Tde1(M) (Fig. 1C, Fig. S1B). This indicates that Tdi1 cannot neutralize the N-terminus-mediated toxicity.

## A glycine zipper motif in N-terminus of Tde1 is required for toxicity and enhanced membrane permeability

To get an insight into the cause of growth inhibition by N-terminus of Tde1, we used N-Tde1 region as a query to search against the NCBI non redundant (nr) database and identified Tde1 homologues encoded in the T6SS gene clusters of *Agrobacterium/Rhizobium* as well as tape measure proteins (TMP) encoded in genomes of *Paraburkholderia/Burkholderia* (Fig. 1D, Fig, S2). We noticed the conservation of continuous copies of GxxxG motifs (12-51) in the N-terminus of Tde1, which resembles the glycine zipper motifs overrepresented in membrane proteins and reported to be involved in the membrane channel formation (<u>Kim</u> *<u>et al.</u>*<u>, 2005</u>). Thus, we hypothesized that these repetitive glycine zipper motifs are involved in membrane permeability and N-Tde1 toxicity.

**Figure 2.**
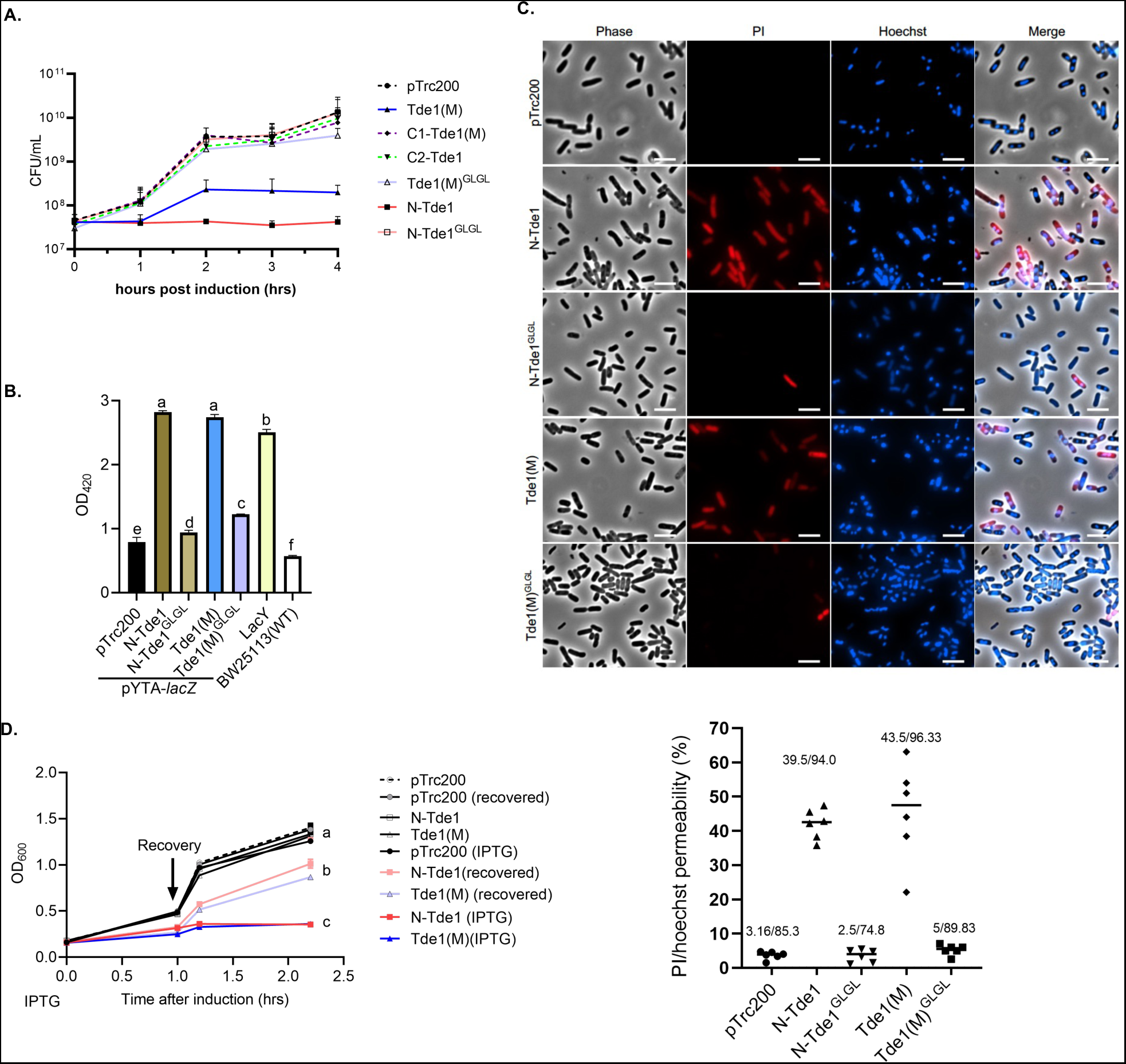
Growth inhibition and membrane permeabilization assays of glycine zipper mutants. (A) Growth inhibition assay of *E. coli* DH10B cells harbouring pTrc200 vector or each of its derivatives expressing Tde1 variants with IPTG inducible expression. The growth of *E. coli* was monitored by CFU counting every 1 hr. For membrane permeabilization assays, BW25113 WT or BW25113 (Δ*lacY,* pYTA-*lacZ)* cells harbouring pTrc200 vector or each of its derivatives expressing Tde1 variants were carried out for (B) β-galactosidase activity assay to determine ONPG uptake, (C) propidium iodide permeability with cells treated with Propidium iodide and Hoechst for detection by fluorescence microscope (Scale bar = 5μm). For quantification of cells with PI signals, a total of 6 randomly selected images obtained from two independent experiments were used to quantify the number of PI stained cells / number of Hoechst stained cells as indicated. (D) Bacteriostatic activity assay. *E. coli* DH10B cells harbouring pTrc200 vector or each of its derivatives expressing Tde1 variants were cultured with or without IPTG induction for 1 hr. The IPTG-induced cells were further centrifuged and resuspended in the fresh medium with or without IPTG. Cell density was measured again before continuous growth for additional 1 hr. Graphs of panels A, B, and D show mean ± SD of three independent experiments. Different letters indicate statistically different groups of strains (p value <0.01).

To verify the hypothesis, two highly conserved glycine residues at position 39 and 43 of a glycine zipper motif were substituted with leucine (G39L and G43L), and the resulting N-Tde1 and Tde1(M) variants were named as N-Tde1^GLGL^, Tde1(M)^GLGL^ respectively. The growth analysis of *E. coli* DH10B cells by counting viable cells and OD_600_ measurement showed that both N-Tde1^GLGL^ and Tde1(M)^GLGL^ lost the ability to cause growth inhibition (Fig. 2A, Fig. S1C). Similar results were also observed when they were overexpressed in *A. tumefaciens Δtde1* mutant (Fig. S1D), indicating that the G^39^xxxG^43^ glycine zipper motif of Tde1 is required for the observed toxicity.

Next, we tested whether N-Tde1 can increase *E. coli* inner membrane permeability. To do so, we used the β-galactosidase activity assay to measure the entry of ortho nitrophenyl galactopyranoside (ONPG) (301 Da) into the cytosol. ONPG normally requires a functional permease LacY to enter into the cytosol but can enter if the inner membrane is permeabilized/compromised (<u>Casteels et al., 1993</u>; <u>Epand et al., 2009</u>). N-Tde1 and Tde1(M) as well as their glycine zipper substitution variants were expressed in *E. coli* BW25113Δ*lacY* (<u>Baba et al., 2006</u>) carrying β-galactosidase (pYTA-*lacZ*). The BW25113Δ*lacY*(pYTA-*lacZ*) complemented with *lacY* was used as a positive control. The *E. coli* cells were induced with IPTG to express Tde1 variants for one hr and collected for ONPG uptake assay. This time point was chosen because there is no obvious difference on the number of viable cells among the strains tested (Fig. 2A, S1E). The results showed that cells expressing either N-Tde1 or Tde1(M) had similar β-galactosidase activity as LacY expressing cells. In contrast, cells expressing N-Tde1^GLGL^ and Tde1(M)^GLGL^ only exhibited background level activity as the negative controls (Fig. 2B). These results indicate that the N-Tde1 and Tde1(M) are able to increase membrane permeability depending on the G^39^xxxG^43^ motif. The data also suggest that the N-terminus-mediated growth inhibition is caused by its ability to enhance inner membrane permeability through glycine zipper motifs.

To further analyse the extent of enhanced membrane permeabilization, cells from the same experiment were normalized to the same OD_600_ and stained with Hoechst and propidium iodide (PI). Hoechst (616 Da) is a nucleic acid staining dye which is permeable to live Gram-negative bacterial cells while PI (668.4 Da) can only enter through a compromised inner membrane or dead cells. The PI/Hoechst staining showed strong PI signals in cells expressing N-Tde1 and Tde1 (M) but no or weak signals were detected in cells expressing N-Tde1^GLGL^, Tde1(M)^GLGL^, or vector control, indicating that N-Tde1 is able to enhance membrane permeability to allow molecules with size 668.4 Da to pass (Fig. 2C).

We next determined whether the N-terminus of Tde1 is bacteriostatic or bacteriolytic by growth recovery assay (<u>Mariano et al., 2019</u>). *E. coli* cells were induced with IPTG to express N-Tde1 or Tde1(M) and after one hr, washed with fresh media without IPTG for continuous cultivation. We found that growth was recovered when cells were washed of the IPTG inducer, in contrast to the growth inhibition of cells with continuous IPTG induction (Fig. 2D). Collectively, the data suggest that the N-terminus of Tde1 is sufficient to facilitate membrane permeability for bacteriostatic toxicity, and such activity requires the conserved G^39^xxxG^43^ glycine zipper motif.

### The N-terminus of Tde1 is necessary and sufficient for Tap1 interaction

Tap1 is the adaptor for loading Tde1 onto VgrG1 (<u>Bondage</u> *<u>et al.</u>*<u>, 2016</u>; <u>Ma</u> *<u>et al.</u>*<u>, 2014</u>). However, the region that Tde1 and Tap1 interact remains undefined. Thus, we performed co-immunoprecipitation (co-IP) assay to identify the specific region of Tde1 that can interact with Tap1 in *A. tumefaciens*. The HA-tagged Tde1 variants were expressed in Δ*tde1* and anti-HA agarose bead was used to co-precipitate the interacting proteins followed by western blotting to detect Tde1 variants and Tap1. The results showed that the N-Tde1 and Tde1(M) interact with Tap1 but not the C-terminal variants, C1-Tde1(M) and C2-Tde1 (Fig. 3A, Fig. S3). N-Tde1^GLGL^ and Tde1(M)^GLGL^ remain capable of interacting with Tap1 (Fig.3A). The results suggest that Tap1 interacts with Tde1 through the N-terminus and that the G39L and G43L substitution does not affect Tde1-Tap1 interaction.

**Figure 3.**
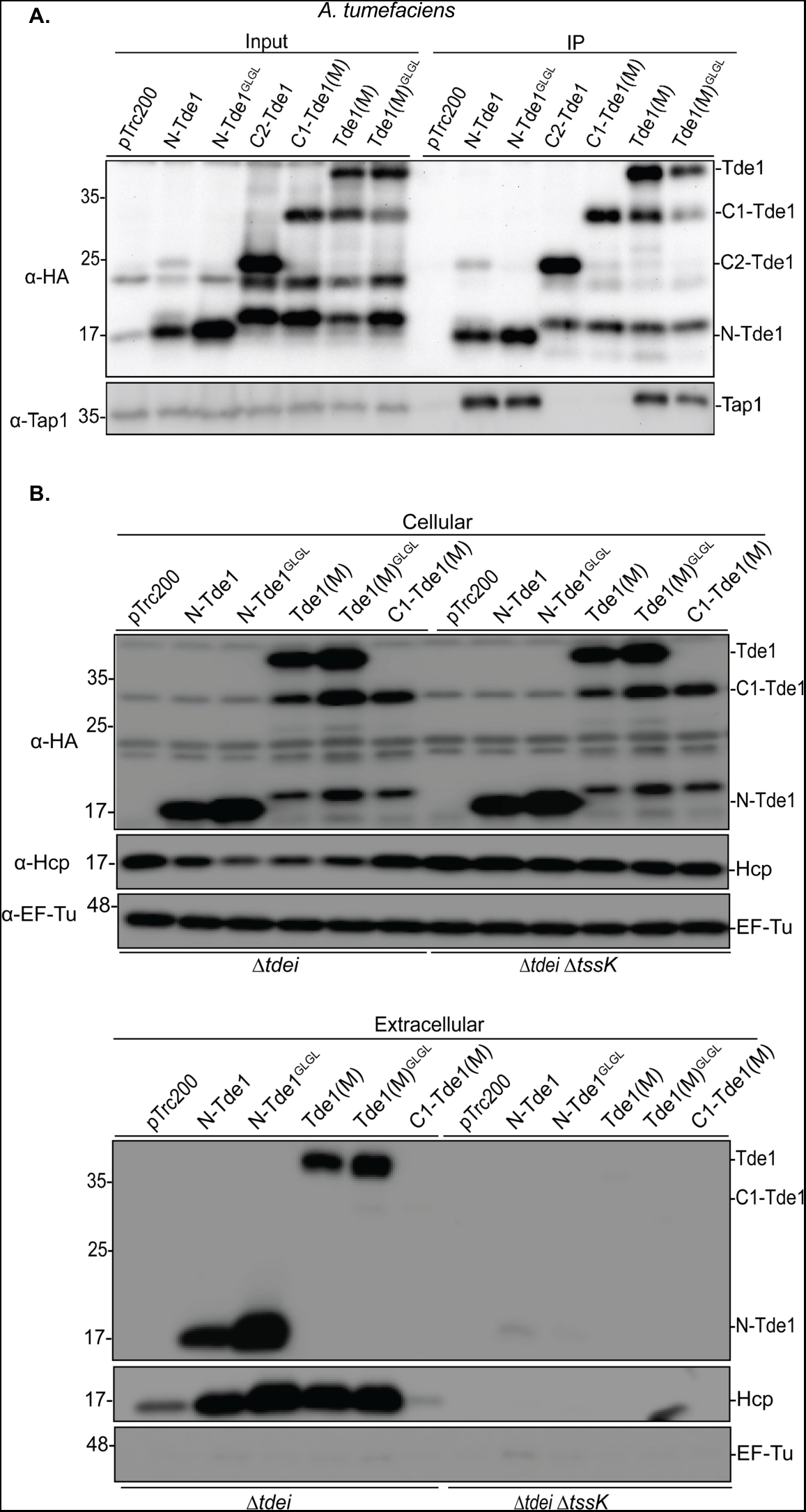
The N-terminus of Tde1 is sufficient for interaction with Tap1 and secretion. (A) Co-immunoprecipitation (Co-IP) in *A. tumefaciens. A. tumefaciens* C58 Δ*tde1* harbouring pTrc200 vector or its derivatives expressing HA-tagged Tde1 variants. Anti-HA resin was used to coprecipitate the Tde1 variants and Tap1. (B) Secretion assay for HA-tagged Tde1 variants. Western blot for the cellular and extracellular fractions of *A. tumefaciens* C58 Δ*tdei* and Δ*tdei1ΔtssK* expressing the HA-tagged Tde1 variants. Hcp secretion was detected as a positive control for active T6SS secretion. Representative western blot results of three independent experiments were shown with antibody against HA, Hcp, or EF-Tu serving as a loading and non-secreted protein control. Protein markers are indicated in kDa.

### The N-terminus of Tde1 is necessary and sufficient for secretion

Because N-Tde1 interacts with Tap1, we hypothesized that this region is required for Tde1 secretion. Thus, we performed secretion assay by expressing the various HA-tagged Tde1 variants in Δ*tdei,* a deletion mutant lacking both *tde1-tdi1* and *tde2-tdi2* toxin immunity pairs. Both cellular and extracellular fractions were collected to determine their expression and secretion, respectively. The results showed that all Tde1 variants containing N-terminus are secreted but not the C-terminus, C1-Tde1(M). The secretion is in a T6SS-dependent manner as secretion was essentially abrogated in Δ*tdeiΔtssK,* which lacks both *tdei* and *tssk* encoding the baseplate component. N-Tde1^GLGL^ and Tde1(M)^GLGL^ are also stably expressed and secreted (Fig. 3B, Fig. S3). The data suggest that N-terminus of Tde1 is necessary and sufficient for secretion and that the G39L and G43L substitution does not interfere with the secretion capacity of Tde1. Accordingly, Hcp secretion levels are highly correlated with Tap1-Tde1 interaction and secretion of Tde1 variants (Fig. 3B). The data also confirmed the requirement of the Tap1-Tde1 interaction for Tde1 secretion and supported our previous finding that Tde loading onto VgrG is critical for active T6SS secretion (<u>Wu et al., 2020</u>).

### G^39^xxxG^43^ motif of Tde1 is required for target cell delivery

Because the G^39^xxxG^43^ glycine zipper motif located in N-Tde1 increased the membrane permeability but was not required for secretion, we hypothesized that G^39^xxxG^43^ is responsible for inserting Tde1 into the inner membrane and delivering it into the cytoplasm of target cells. Here, we engineered each of Tde1 variants fused to super-folder green fluorescence protein (sfGFP) with a flexible (GGGS) linker between Tde1 and sfGFP to avoid the Tde1 functional/structural interference by GFP. The sfGFP fused Tde1 variants were expressed in *A. tumefaciens Δtdei* and Δ*tdeiΔtssK* mutants. The secretion assay results showed that both WT and G39L and G43L substitution of N-Tde1-sfGFP and Tde1(M)-sfGFP are secreted (Fig. S4A). No or trace amounts of proteins were observed in the extracellular fractions of Δ*tdei1Δtssk* mutants, demonstrating that the secretion was T6SS dependent. C1-Tde1(M)-sfGFP protein signal could not be unambiguously determined in the cellular fraction due to the overlapping of its predicted protein band with cross-reacted proteins, and no corresponding C1-Tde1(M)-sfGFP band was detected in the extracellular fraction. The secretion assay of Tde1 variants fused with either HA or sfGFP concluded that N-Tde1 is necessary and sufficient for secretion and the G39L and G43L substitution does not affect Tde1 being secreted, which is correlated with the ability to interact with Tap1.

We next investigated the translocation of Tde1 variants by mixing *A. tumefaciens Δtdei,* expressing sfGFP-fused Tde1 variants, with *E. coli* cells expressing mCherry. *A. tumefaciens* expressing sfGFP only (Vector-sfGFP) was used as a negative control. After co-culture, we imaged populations for mCherry (false coloured in blue) and GFP (green) to detect *E. coli* and *A. tumefaciens*, respectively. We merged images to identify cyan colored cells (overlayed blue and green signals), which represent *E. coli* cells with translocated Tde1 variants carrying sfGFP. We were able to observe ∼50% of cells with cyan fluorescence when *A. tumefaciens* expressing N-Tde1-sfGFP and Tde1(M)-sfGFP was co-cultured with *E. coli* mCherry whereas the GFP and mCherry signals were not overlapped in the *E. coli* cells co-cultured with *A. tumefaciens* strains expressing GFP only or sfGFP-fused C1-Tde1(M), N-Tde1^GLGL^, Tde1(M)^GLGL^ respectively (Fig. 4). No cyan fluorescence was observed when N-Tde1-sfGFP and Tde1(M)-sfGFP were expressed in the Δ*tdei1Δtssk* mutant as the attacker (Fig. S4B).

**Figure 4.**
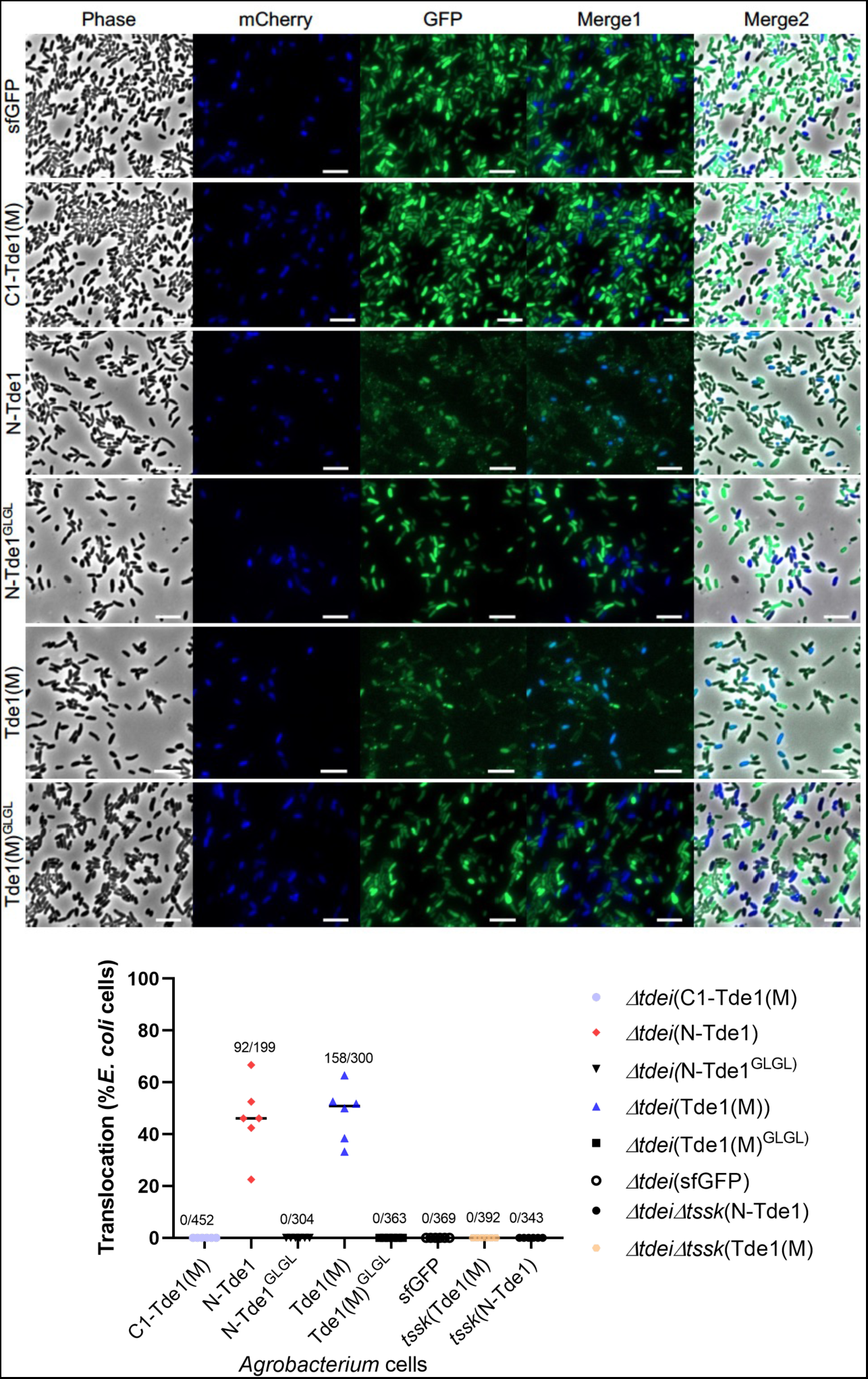
Translocation of Tde1 variants fused with sfGFP by *A. tumefaciens-E. coli* co-culture. Fluorescence microscopy for Tde1 translocation. *A. tumefaciens* C58 Δ*tdei* expressing Tde1 variants fused with sfGFP (in green) and *E. coli* DH10B carrying mCherry (false coloured in blue) were co-cultured for 20 hrs. A cyan fluorescence with merged blue and green signals represented the translocation of Tde1 variants from *A. tumefaciens* to *E. coli* (Scale bar = 5μm). The number of cells with overlayed GFP and mCherry fluorescence was quantified from a total of 6 randomly selected images obtained from three independent experiments (number of cells with cyan fluorescence/ total *E. coli* cells counted).

The data suggest that Tde1 is translocated into target cells in a T6SS- and G^39^xxxG^43^- dependent manner. Because N-Tde1^GLGL^-sfGFP and Tde1(M)^GLGL^-sfGFP could be secreted but not translocated into target cells, G^39^xxxG^43^ motif is necessary for delivering Tde1 into the target cell.

### G^39^xxxG^43^ is critical for interbacterial competition but not for DNase activity

To assess the role of the G^39^xxxG^43^ motif for target cell intoxication in the context of interbacterial competition, *A. tumefaciens* C58 Δ*tdei* expressing either Tde1-Tdi1, Tde1(M)-Tdi1, or single/double G39L and G43L substitution variants, was competed with target *E. coli* (DH10B) cells. By counting the survival rate of *E. coli* prey cells, the data showed that *A. tumefaciens Δtdei* (Tde1-Tdei1) exhibits an antibacterial activity but not in the negative controls, the secretion deficient mutants Δ*tssL* and Δ*tdei*Δ*tssK* (Tde1-Tdei1) (Fig. 5A). No antibacterial activity could be observed for *A. tumefaciens Δtdei* expressing Tde1(M)-Tdi1, indicating the DNase-mediated killing to *E. coli*. The antibacterial activity of Δ*tdei*(Tde1^GLGL^-Tdi1, Tde1^G39L^-Tdi1, Tde1^G43L^-Tdi1) was not detectable, similar to that of negative controls. We also performed interbacterial competition assays using *A. tumefaciens* strain 1D1609, which is susceptible to T6SS killing by C58 (<u>Wu et al., 2019</u>). The interbacterial competition between two *A. tumefaciens* strains was calculated by competitive index, which revealed the higher competitiveness of Δ*tdei* (Tde1-Tdi1) and C58 against 1D1609 but no competitive advantage could be detectable for any of glycine zipper variants or Tde1(M) (Fig. 5B). The observed antibacterial activity is T6SS-dependent because the killing activity of Tde1 was not observed when expressed in Δ*tdeiΔtssK.* The results indicate that G^39^xxxG^43^ motif is required for interbacterial competition at both inter- or intra-species levels. We also performed a secretion assay of these *A. tumefaciens* attacker strains and all glycine zipper variants were secreted (Fig. 5C). It is notable that Tde1^GLGL^ proteins accumulated at slightly lower levels while Tde1^G39L^ and Tde1^G43L^ had similar or even higher protein levels to that of Tde1 and Tde1(M). Accordingly, Tde1^GLGL^ was secreted at lower levels.

**Figure 5.**
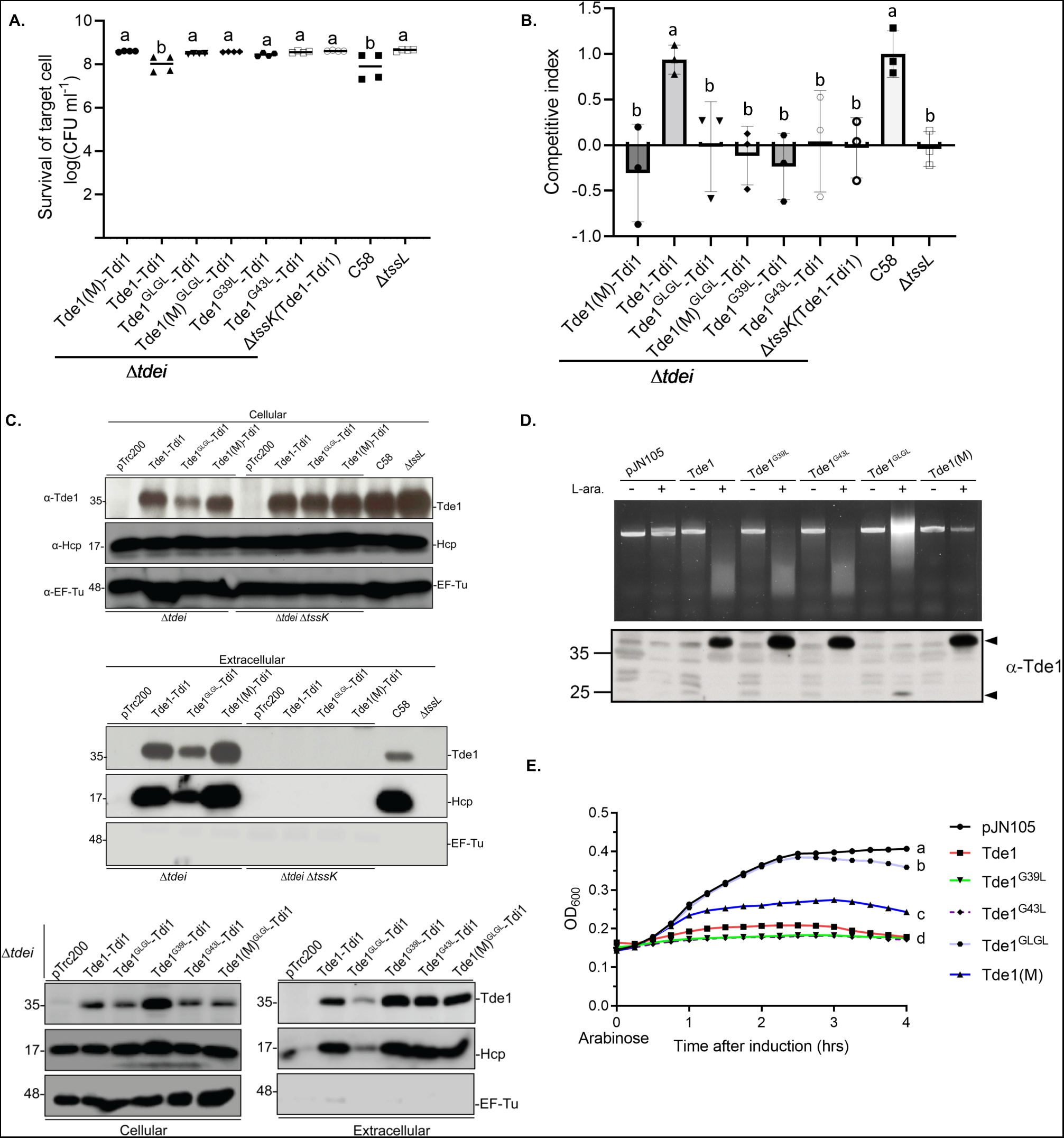
G^39^xxxG^43^ glycine zipper motif of Tde1 is required for DNase-mediated killing of target cells during interbacterial competition. (A) Interbacterial competition of *A. tumefaciens* C58 Δ*tdei* and Δ*tdei1ΔtssK* expressing the Tde1 variants against *E. coli* cells was carried out on LB medium and *E. coli* survival rate was quantified by CFU counting. (B) Interbacterial competition between various *A. tumefaciens* C58 strains and *A. tumefaciens* 1D1609 on AK medium and the competition outcome was shown by competitive index. (C) Secretion assay for Tde1 and its variants co-expressed with its immunity protein Tdi1 in *A. tumefaciens* C58 Δ*tdei* and Δ*tdeiΔtssK.* (D) *In vivo* plasmid DNA degradation assay. *E. coli* BW25113 carrying pJN105 empty vector or the derivatives expressing different variants of Tde1 were supplemented with 0.5% glucose (“-”) or 0.2% L-arabinose (“+”) for 3 hrs to either repress or induce Tde1 production. The plasmids were then extracted to observe the DNA degradation and bottom panel showed western blots of specific Tde1 protein bands indicated by arrows. (E) Growth inhibition assay of Tde1 and its variants. *E. coli* BW25113 cells were induced by adding 0.2% L-arabinose for Tde1 production. The OD_600_ values were measured every 15 minutes. The OD_600_ values of the 4 h post L-arabinose induction were analysed for statistical analysis. Graphs show mean ± SD of three biological repeats. Western blots were detected with specific antibody against Tde1, Hcp, or EF-Tu serving as a loading and non-secreted protein control. Protein markers are indicated in kDa. Data in panel A are mean ± SD of four biological repeats of two independent experiments. Panels B and E show mean ± SD of three independent experiments. Different letters indicate statistically different groups of strains (p value <0.05). Results in panel C and D are representative of three independent experiments.

To exclude the possibility that G39L and G43L substitution may influence its DNase activity, we performed *in vivo* plasmid DNA degradation assay. Tde1 and the variants were each expressed by tightly controlled arabinose-inducible promoter for *in vivo* plasmid DNA degradation assay in *E. coli* BW25113 as described (<u>Ma *et al*., 2014</u>). It was observed that plasmid DNA was completely degraded in cells expressing Tde1, but not in the negative controls, the cells without arabinose induction nor cells expressing Tde1(M). Plasmid DNA was also degraded by Tde1^GLGL^ but not as complete as Tde1 while both Tde1^G39L^ and Tde1^G43L^ exhibit wild-type level DNase activity. (Fig. 5D). The lower DNA degradation efficiency of Tde1^GLGL^ could be correlated with the barely detected Tde1^GLGL^ proteins (Fig. 5D). We also found that the degree of plasmd DNA degration is also correlated with the growth inhibition effect (Fig. 5E and Fig. S1E). The slightly recovery of Tde1^GLGL^ from growth inhibition is consistent with the instability of Tde1^GLGL^. The evidence that G39L and G43L substitutions abolished interbacterial competition but did not affect DNase activity and secretion of Tde1 suggest the G^39^xxxG^43^ motif is necessary for delivering Tde1 across the inner membrane into the cytoplasm of target cells.

## DISCUSSION

Through the dissection of Tde1 DNase effector, we provide strong evidence for a role of the N-terminal glycine-zipper motif(s) of Tde1 in delivering the T6SS effector into target cells. Here, we propose a model explaining the loading, firing, and translocation of Tde1 (Fig.6). In *A. tumefaciens,* Tde1 DNase activity is neutralized by Tdi1 by binding to C-terminal DNase domain while its N-terminal domain interacts with Tap1 for loading onto VgrG1 (Step 1). The VgrG1-Tap1-Tde1-Tdi1 complex is then recruited onto membrane-associated baseplate, which serves as a docking site for polymerization of Hcp tube and TssBC sheath (Step 2). Upon TssBC sheath contraction (Step 3), Tap1 and Tdi1 may fall off and Hcp-VgrG-Tde1 puncturing device is then ejected for secretion. In contact with a target cell, Tde1 may be delivered to periplasm of the target cell where Tde1 permeabilises the inner membrane in a G^39^xxxG^43^ motif-dependent manner (Step 5). Once delivered, Tde1 exerts its toxicity by attacking DNA for degradation (Step 6).

**Figure 6.**
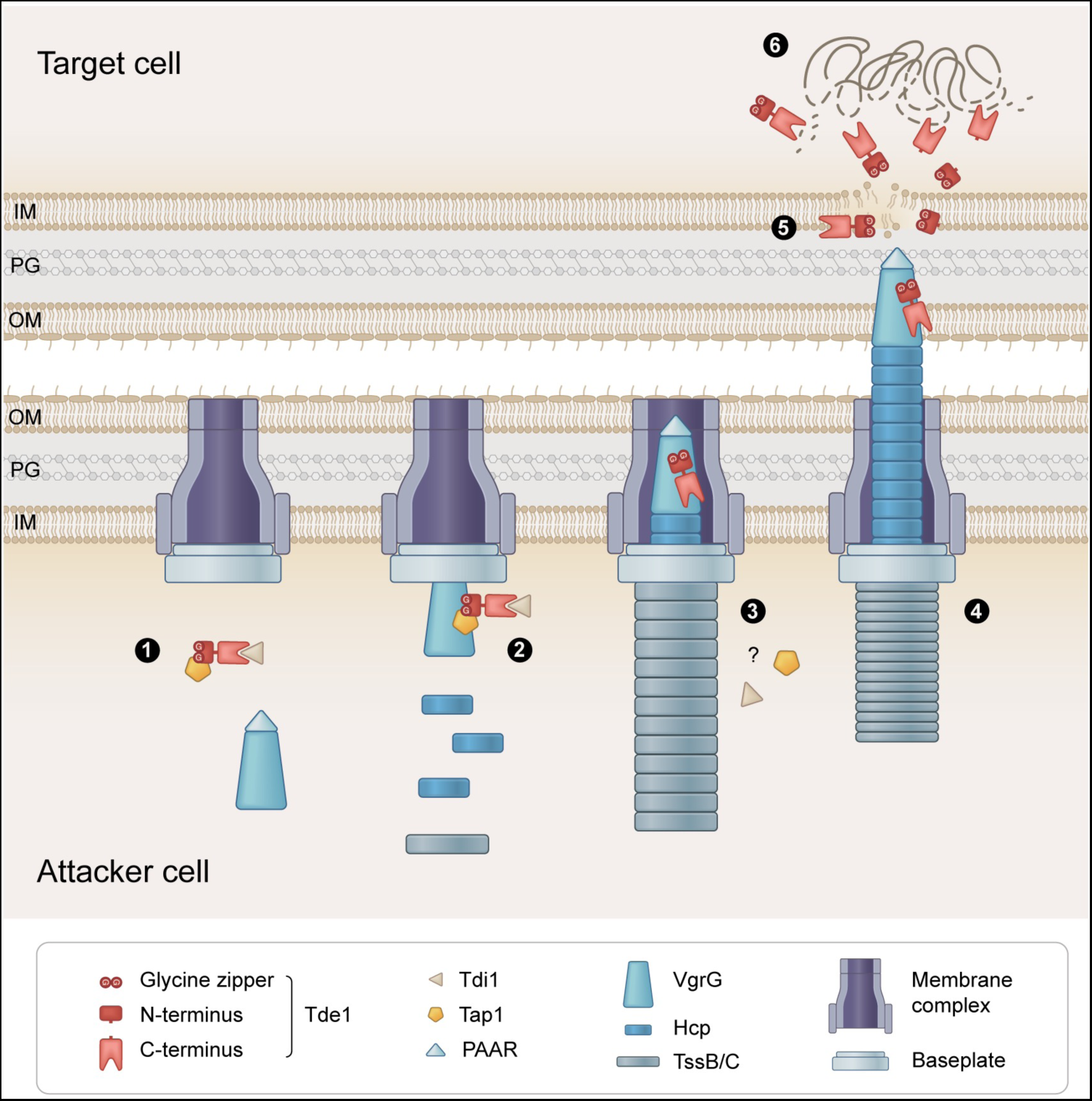
Proposed model of the loading, firing, and translocation of Tde1. The Tde1 translocation is proposed through six steps. Step 1: Tde1 forms a complex with Tdi1 and Tap1 in the attacker cell. Step 2: Tap1-Tde1-Tdi1 complex binds to the VgrG and the Hcp-VgrG-PAAR puncturing device carrying Tde1 effector complex is loaded onto membrane-associated baseplate. Step 3: Hcp tube and TssB/C sheath polymerize on the Tde1-loaded VgrG/baseplate while Tdi1 and Tap1 fall off with unknown mechanisms before or upon firing. Step 4: TssBC sheath contracts and ejects Tde1 into the target cell periplasm or cytoplasm. Step 5: The glycine zipper(s) on the N-terminus of Tde1 permeabilize the target cell membrane. Step 6: Intact or truncated Tde1 proteins attack DNA for degradation in the target cell.

T6SS cargo effectors often require the specific chaperone/adaptor for loading onto puncturing device for secretion. Our previous findings demonstrated that Tap1, a DUF4123-containing protein, specifically interacts with Tde1 for loading onto VgrG1 for secretion (<u>Bondage</u> *<u>et al.</u>*<u>, 2016</u>; <u>Ma</u> *<u>et al.</u>*<u>, 2014</u>). We now show that N-terminal region of Tde1 is necessary and sufficient for interaction with Tap1 for secretion and delivery into target cells. The evidence that Tde1^GLGL^ variant remains capable for binding to Tap1 for export but deficient in membrane permeability and translocation demonstrate a distinct role of this G^39^xxxG^43^ motif in target cell delivery. Among the nine classes of the Ntox15-containing proteins, the majority of them including Tde1 belong to class I without detectable N-terminal domains (Fig. S2). We identified the presence of glycine zipper motifs overlapping with transmembrane domain (TMD) not only in N-terminal region of all Tde1 orthologs encoded in Rhizobiaceae but also in C-terminal region of tape measure proteins (TMP) encoded in genomes of *Paraburkholderia/Burkholderia,* likely as a prophage. TMP is a phage protein suggested to have a channel forming activity (<u>Roessner and Ihler, 1984</u>; <u>1986</u>) and as a determinant in connecting host inner membrane proteins for injecting phage genome into bacterial host cells (<u>Cumby et al., 2015</u>). Such conservation in Tde1 orthologs suggests that this glycine zipper-mediated delivery could be a common strategy deployed by these bacterial effectors for translocation across target cell membranes. It would be also interesting to investigate whether TMP also employs its C-terminal glycine zipper to mediate phage genome entry into host cells.

A role of N-terminal domain involved in translocation of polymorphic toxins have been well documented in those of contact-dependent growth inhibition (CDI) system and bacteriocins (<u>Ruhe et al., 2020</u>). However, little is known for the translocation mode of bacterial toxins delivered by other systems. Previous study in *P. aeruginosa* showed that VgrG-loaded Tse6-EgaT6 complex is sufficient to translocate across a lipid bilayer *in vitro* (<u>Quentin et al., 2018</u>), suggesting a role of VgrG-effector complex itself in inserting across the inner membrane of target cells. A recent study further uncovered a widespread prePAAR motif in N-terminal TMDs of T6SS effectors involved in interaction with Eag family chaperone for export (<u>Ahmad et al., 2020</u>). The findings from the Tap1 and Eag chaperone-mediated T6SS toxins led us to propose that the bacterial toxins harbouring a N-terminal TMD may be protected by its cognate chaperone/adaptor from insertion into membranes in the attacker cell. However, once the effector is injected into the periplasm of the target cell, specific motifs (such as glycine zippers or perhaps prePAAR) may insert into the inner membrane for the delivery into the cytoplasm. By an elegant *in vitro* translocation assay, a recent study discovered a N-terminal domain of a bacteriocin pyocin G is required for import of its C-terminal nuclease toxin into the cytoplasm cross inner membrane (<u>Atanaskovic et al., 2022</u>). This inner membrane translocation domain (IMD) is distinct from the glycine zipper repeats identified in this study but also found conserved in other bacterial toxins including some of T6SS. Thus, bacterial toxins itself directing translocation into target cells could be a general strategy used by bacteria for interbacterial competition.

A few membrane permeabilizing T6SS toxins have been reported. The *Vibrio cholerae* VasX caused dissipation of membrane potential, leading to membrane permeabilization of target bacterial cells similar to the Tme effectors of *V. parahaemolyticus,* which represents a widespread family of T6SS effectors harbouring C-terminal TMD for membrane disruption (<u>Fridman et al., 2020</u>; <u>Miyata et al., 2013</u>). On the other hand, Tse4 disrupts the membrane potential and forms a cation selective pore without membrane permeabilization where the pore cannot even allow the permeability of a relatively smaller molecular weight (ONPG, 300 Da) (<u>LaCourse</u> *<u>et al.</u>*<u>, 2018</u>). Distinct from these toxins in which they confer pore forming activity for toxicity, the role of glycine zipper(s) of Tde1 appears to enhance membrane permeability for bringing the toxin domain into target cell cytoplasm because Tde1(M) with complete glycine zipper motifs but loss of DNase activity did not show interbacterial competition activity against *E. coli* or *A. tumefaciens* under conditions tested (Fig. 5A, 5B) (<u>Ma</u> *<u>et al.</u>*<u>, 2014</u>).

To date, no structural information is available for Ntox15 superfamily proteins where Tde1 belongs. While N-terminus of Tde1 lacks sequence similarity to any of those known pore-forming toxins, structural modelling showed structural similarity of two helixes containing consecutive glycine zipper motifs to the pore forming domain of pyocin S5 (<u>Behrens et al., 2020</u>) (Fig.S6). Pyocin S5 can cause ATP leakage and PI permeability (<u>Ling et al., 2010</u>) potentially to inner membrane after translocation through FptA and TonB1 (<u>Behrens *et al*., 2020</u>). Tde1 allows the passage of relatively larger molecule, PI (668 Da), suggesting that its N-terminal glycine zipper(s) may form larger pores similar to pyocin S5. G^39^xxxG^43^ motif plays no role in DNase activity of Tde1 but is crucial for its protein stability. Tde1 with substitution of one of the two glycine residues to leucine retains stability of intact proteins but Tde1 is prone for truncations and degradation when both glycine residues are substituted to leucine. The instability is most evident when ectopically expressed in *E. coli* and when retaining DNase activity (Fig. S1A, Fig. 5). Single glycine substitution (Tde1^G39L^ and Tde1^G43L^ variants) does not influence protein stability may suggest that the adjacent glycine residues (G^35^ or G^47^) are sufficient to compensate the loss of one glycine of G^39^xxxG^43^ motif for structural integrity in both variants. The importance of G^39^xxxG^43^ motif in Tde1 protein stability is consistent with the role of glycine zippers in structural impact (<u>Kim *et al*., 2005</u>). However, integrity of G^39^xxxG^43^ motif is critical for interbacterial competition because both Tde1^G39L^ and Tde1^G43L^ variants do not exhibit detectable antibacterial activity to either *E. coli or A. tumefaciens* (Fig. 5A, 5B). These results suggest the role of G^39^xxxG^43^ motif in delivering Tde1 across the inner membrane into the cytoplasm of target cells.

It is striking to observe such a high percentage of cells (∼50%) representing N-Tde1-sfGFP and Tde1(M)-sfGFP translocation from *A. tumefaciens* into *E. coli* cells expressing mCherry (Fig. 4). Adding the flexible GGGS linker between sfGFP and Tde1 that retain both Tde1 secretion activity and GFP fluorescence may be the key for the success of this translocation experiment. Interestingly, we also observed many GFP foci from *A. tumefaciens* cells expressing translocation competent N-Tde1-sfGFP or Tde1(M)-sfGFP while others including *E. coli* cells with GFP signals uniformly distributed throughout the cells. Based on the role of glycine zippers in interacting with membrane, we propose that Tde1 proteins may preferentially bind to microdomain of cytoplasmic membrane, which was recently found in *A. tumefaciens* (<u>Czolkoss et al., 2021</u>). We also found that Tde1 proteins (either tagged with HA or GFP, Fig. 3, S1, S4A) are prone for truncation especially when they are ectopically expressed in *E. coli* or when Tdi1 is absent or not equivalent. Thus, it is possible that Tde1-GFP proteins are truncated after translocation into *E. coli* cells, in which most GFP signals are emitted from free GFP instead of Tde1-GFP. The stability of free GFP derived from translocated Tde1-GFP may also explain the high percentage of *E. coli* cells exhibiting overlayed GFP/mCherry signals. There is evidence that truncation of T6SS effectors is critical for the toxicity (<u>Pei et al., 2020</u>). Future work to investigate how Tde1 interacts with membrane and dissect the region required for DNase activity shall shed light to understand the biological significance and mechanisms underlying this interesting observation.

With the knowledge of effector translocation mechanisms, the bacterial protein secretion apparatus also offers a strategy for delivering heterologous proteins to specific cells. T6SS is a promising vehicle for such purpose because effectors or secreted proteins appear to be delivered with their folded or partially folded form, unlike those to be transported as unfolded forms in most of other specialized secretion systems (<u>Costa *et al*., 2015</u>). Engineering T6SS carriers such as VgrG spikes to carry exogenous effectors proteins into target cells are feasible but challenging (<u>Ho et al., 2017</u>; <u>Wettstadt and Filloux, 2020</u>; <u>Wettstadt et al., 2020</u>). By using a truncated variant of PAAR, a recent study showed delivering exogenous T6SS effectors and Cre recombinase for genetic modification in the target cells (<u>Hersch et al., 2021</u>). Our success in using N-Tde1 in the delivery of sfGFP proteins into target *E. coli* cells where they exert fluorescence also suggest potential applications of N-Tde1 for the delivery of proteins of interests such as genetic modifiers. This strategy provides advantages over transforming foreign DNA for expressing a protein of interest from creating undesired genome manipulation.

## MATERIALS AND METHODS

### Strains and growth conditions

The strains and plasmids used in this study are listed in the supplementary Tables S1 and S2. The *E. coli* strains used in this study are BW25113 and DH10B. All the *A. tumefaciens* strains were cultured on 523 medium (<u>Kado, 1970</u>) at 28 °C unless stated. The *E. coli* strains were cultured on Luria Bertani (LB) medium (10 g L−1 NaCl, 10 g L−1 tryptone, and 5 g L−1 yeast extract) at 37°C unless stated. Where appropriate, the media were supplemented with 100 μg mL^−1^ spectinomycin, gentamycin 25 μg mL^−1^ (for *E. coli*) and 50 μg mL^−1^ (for *A. tumefaciens*), 50 μg mL^−1^ ampicillin, 50 μg mL^−1^ kanamycin, 1 mM Isopropyl β- d-1-thiogalactopyranoside (IPTG).

### Growth inhibition assay

For growth inhibition assay of IPTG-inducible expression of Tde1 and its variants, *E. coli* (DH10B) harbouring pTrc200 vector or the derivatives expressing Tde1 variants were grown overnight in LB medium supplemented with spectinomycin prior to 1:30 dilution in a fresh medium and incubated for 2 hrs at 37°C with 250 rpm. After 2 hrs, the cultures were normalized to OD_600_ 0.1 in LB with 1 mM IPTG for continuous culture in the same growth condition. The growth of *E. coli* was monitored for OD_600_ every one hr using ULTROSPEC^®^ 10 cell density meter (Biochrom, UK) or viable cell by counting colony forming units (CFUs) on LB agar containing Sp. For growth inhibition assay of arabinose-inducible expression, *E. coli* BW25113 harbouring pJN105 vector or the derivatives expressing Tde1 variants were used. Overnight cultures of *E. coli* cells were adjusted to an OD_600_ of 0.1 in 200 μL LB with 0.2% L-arabinose in a 96-well plate. The OD_600_ values were measured by the Synergy H1 microplate reader (Agilent Technologies, USA) with agitation at 37 °C. The OD_600_ values or CFUs of indicated time points were used to calculate mean ± SD of three independent repeats. One way Analysis of Variance (ANOVA) was used for the analysis of statistical significance followed by the Tukey’s multiple comparison.

#### *In vivo* plasmid DNA degradation assay

The *in vivo* plasmid DNA degradation assay was performed as described (<u>Ma *et al*., 2014</u>) with minor modifications. Briefly, overnight cultures of *E. coli* BW25113 carrying pJN105 vector or the derivatives expressing Tde1 variants were adjusted to an OD_600_ of 0.3 in 4 mL LB with 0.5% D-glucose or 0.2% L-arabinose. After induction for 3 hrs, bacterial cells normalized by OD_600_ were collected for plasmid DNA extraction and western blot analysis. The plasmids were then extracted and applied to 0.6% agarose gel electrophoresis to detect the DNA degradation. The OD_600_ values were measured by DEN-600 photometer (Biosan, Latvia) every hr.

### β-galactosidase and viability assays for ONPG update

β-galactosidase assay was performed as described (<u>Saint Jean et al., 2018</u>) with minor modifications. BW25113 wild type, BW25113Δ*lacY*(pYTA-lacZ), or BW25113Δ*lacY* harbouring pTrc200 vector or the derivatives expressing Tde1 variants were grown overnight and refreshed to a fresh medium as stated for growth inhibition assay. After subculture for 2 hr, the cells were induced with 1 mM IPTG, and incubated for one more hr. Part of the culture was adjusted to OD_600_ = 0.3 in Z-buffer and the Intracellular β-galactosidase activity was measured by mixing 100 µL of 4 mg/ml ONPG with 900 µL of the cells and incubation at room temperature for 10 min prior to measurement at OD_420_. The remaining cells were normalized to OD_600_ 0.3 in 0.9 % sterile saline and after serial dilution, 10 µL was spotted on LB plate without antibiotics to recover the viable cells. Data of OD_420_ were used to calculate mean ± SD of three independent experiments. One way ANOVA was used for the analysis of statistical significance followed by the Tukey’s multiple comparison.

### Co-immunoprecipitation (Co-IP)

The co-IP was performed according to the manufacturer’s recommendations of EZview red Anti-HA agarose (Sigma-E6779) with minor modifications. To identify Tap1 interacting domain of Tde1, the HA tagged Tde1 variants were expressed on pTrc200 plasmid. For co-IP in *A. tumefaciens,* C58 Δ*tde1* cells expressing the Tde1 variants grown in 523 medium overnight were resuspended in a 1:30 ratio to a fresh medium and incubated at 25°C for 3 hrs followed by 1 mM IPTG induction for additional 3 hrs. After 6 hrs post incubation, the cells were normalized to OD_600_ of 5 per mL in ice-cold PBS buffer (pH 7.4). After cell lysis by lysozyme treatment and sonication, the lysate was centrifuged and a 100 μL aliquot of the lysate was saved for the input fraction. The remaining 900 µL lysate was mixed with 25 µL of pre-equilibrated Ezview red Anti-HA agarose and incubated at 4 °C for 1 hr. The beads were then washed 3 times with ice-cold PBS buffer and the proteins bound to the beads were eluted with 100 µL of 2X SDS sample loading buffer. Similarly, the aliquoted input fraction was mixed with equal volume of 2X SDS sample loading buffer for analysis by western blotting.

### Secretion assay

Type VI secretion assay was performed in 523 as described previously (<u>Wu *et al*., 2020</u>). Briefly, *A. tumefaciens* strain was cultured overnight in 523 medium and normalized to OD_600_ of 0.2 in a fresh medium. After 6 hrs of culturing, the secreted proteins were collected by centrifuging at 10,000 g for 5 mins. The resulting pellet was adjusted to OD_600_ of 10 as a cellular fraction. The culture supernatant was filtered with 0.22 μm Millipore filter membrane and resulting filtrate was subjected for TCA precipitation (<u>Wu et al., 2008</u>) and referred as an extracellular fraction.

### Western blotting

Western blot analyses were done as previously described (<u>Lin et al., 2013</u>). The following primary antibody titres used were: HA epitope (1:4,000), Tap1 (1:3,000) (<u>Lin *et al*., 2013</u>), Strep (1:4,000), EF-Tu (1:6,000), and C-terminal Tde1 (1:4,000) (<u>Bondage *et al*., 2016</u>; <u>Ma *et al*., 2014</u>), Hcp (1:2,500) (<u>Wu *et al*., 2008</u>).

### Interbacterial competition assays

For interbacterial competition with *E. coli* (target), *A. tumefaciens* strain (attacker) was grown overnight at 28°C in 523 broth with appropriate antibiotics if needed. *E. coli* DH10B harbouring pRL662 plasmid was grown at 37°C in LB broth with gentamycin. After harvesting and washing the cells in 0.9% saline, the attacker to target cell density was adjusted to 30:1 (OD_600_ =3: 0.1) and the mix was spotted on Agrobacterium kill-triggering (AK) (<u>Yu *et al*., 2020</u>) or LB medium containing 1.5% (wt/vol) agar. After incubation of the mixed strains for 16 hr at 28°C, the spot was resuspended in 0.9% saline, serial diluted, and spotted on a gentamycin-containing LB agar square plate at 37°C to calculate *E. coli* survival rate by CFU counts. Similar procedure was used when using *A. tumefaciens* strain 1D1609 as a target, which was grown at 28°C in 523 broth prior to competition. The competition was carried out on AK medium for 16 hr at 28°C with CFU counting at both initial and final time points by selection of C58 strains with Sp resistance and 1D1609 with Gm resistance. To calculate competitive index, CFUs of *A. tumefaciens* attacker C58 strain were divided by the CFUs of target 1D1609 strain at both 0 hr and 16 hr; and the ratio of C58/1D1609 at 16 hr was divided by the ratio of C58/1D1609 at 0 hr to obtain competitive index. One way Analysis of Variance (ANOVA) was used for the analysis of statistical significance followed by the Fishers Least Significant Difference (LSD) test.

### Fluorescence microscopy

For propidium iodide and Hoechst staining, *E. coli* cells (BW25113) harbouring Trc200 vector or derivatives expressing Tde1 variants were grown overnight and refreshed to a fresh medium as stated for growth inhibition assay. After subculture for 2 hrs, the cells were induced with 1 mM IPTG for one hr and OD_600_ equivalent to 0.3 was collected in 1 ml PBS and stained with Hoechst 33342 (H3570) to a final concentration of 12.3 µg/mL and 1 µg/ml Propidium iodide (2208511) and incubated for 2 minutes in dark. Finally, 2 µL was spotted on 2.5% agarose pad.

For the translocation experiment, the sfGFP fused Tde1 variants were expressed in *A. tumefaciens Δtdei* cells (attacker). *E. coli* (target) cells were labelled with mCherry (false colour blue) expressed from pBBRMCS2. *A. tumefaciens* attacker cells were cultured in 523 broth overnight, and *E. coli* target cells were separately cultured on LB broth. Overnight cultured attacker and target cells were mixed at 5:1 ratio (OD_600_= 1.0: 0.2), and 10 μL of the mix was cultured on LB-agar plate without IPTG. After 20 hrs of co-culture, the cells were washed with 100 μL PBS and 2 µL of suspension was spotted on the 2.5% agarose pad on a microscopic slide. The merged GFP and mCherry signals was based on a cyan fluorescence.

Fluorescence microscopy was performed using Axio Observer 7 (Zeiss, Germany) microscope equipped with an Axiochem 702 digital camera and a Plan-Apochromat 100x/1.4 Oil DIC H objective. Exposure times were adjusted to 20 ms for Phase, 50 ms for Hoechst, 150 ms for PI, 200 ms for GFP and 5000 ms for mCherry. Multiple images were taken from different fields and all the experiments were performed at least in triplicate and a representative image is shown. Images were analyzed by using ZEN 2.3 (blue edition) software.

### Domain prediction and analysis

Full-length Tde1 (1-278) was used as a query for conserved domain search on the conserved domains database (CDD) (<u>Lu et al., 2020</u>) of the National Center for Biotechnology Information (NCBI). Prediction of transmembrane domain was done using the PRED-TMR2 (<u>Pasquier et al., 1999</u>). The Tde1 homologues and tape measure proteins (TMPs) for the multiple sequence alignment were obtained by BLAST search of N-Tde1 (1-97) against the NCBI non redundant database (nr) with representative sequences selected for multiple sequence alignment. The domain architectures of the Ntox15 domain containing proteins were obtained using the full length Tde1 against Conserved Domain Architectural retrieval tool (CDART) of NCBI. The information of gene clusters encoding Tde1 homologues and TMPs including upstream and downstream three genes was retrieved from their respective genomes. Full-length Tde1 (1-278) was used as a query for structural prediction on a Phyre2 (<u>Kelley et al., 2015</u>). Three-dimensional structure modelling was done using Phyre2 in intensive modelling mode. Crystal structure served as best template for the N-terminus and percentage of confidence for three-dimensional structure modelling are indicated in the legends of corresponding figures. The structural graphics were generated by using ChimeraX 1.1 (<u>Goddard et al., 2018</u>).

## ACKNOWLEDGEMENT

We would like to thank the Lai lab members for their help and fruitful discussion throughout this study, Yun-Wei Lien for generating the Δ*tdeiΔtssK* strain, and Chih-Horng Kuo for discussion and suggestion on BLAST analysis. The authors highly appreciate Jeff Chang, See-Yeun Ting, Lay-Sun Ma, and Chih-Feng Wu for critically reading the manuscript and their valuable comments. We also acknowledge the technical assistance of fluorescence microscope provided by Mei-Jane Fang from Live Cell Imaging Division of Cell Biology Core and Sanger DNA sequencing service provided by Genomic Technology Core, both located at the Institute of Plant and Microbial Biology, Academia Sinica. The authors also thank Ying Wang for illustration of proposed model. The funding was supported by National Science and Technology Council (NSTC) of Taiwan (NSTC 110-2311-B-001-032-MY3) and Academia Sinica Investigator Award (AS-IA-107-L01) to EML. YWC was supported by the postdoctoral fellowship (NSCT 110-2811-B-001-524). The funders had no role in study design, data collection and interpretation, or the decision to submit the work for publication.

## AUTHOR CONTRIBUTION

Conceptualization: JA, MY, EML; Investigation: JA, MY, LKS, YWC; Project Funding acquisition and administration: EML; Supervision: EML; Writing of original draft: JA, EML; Writing – methodology, review & editing: JA, MY, LKS, YWC, EML.

---

**Figure S1.**
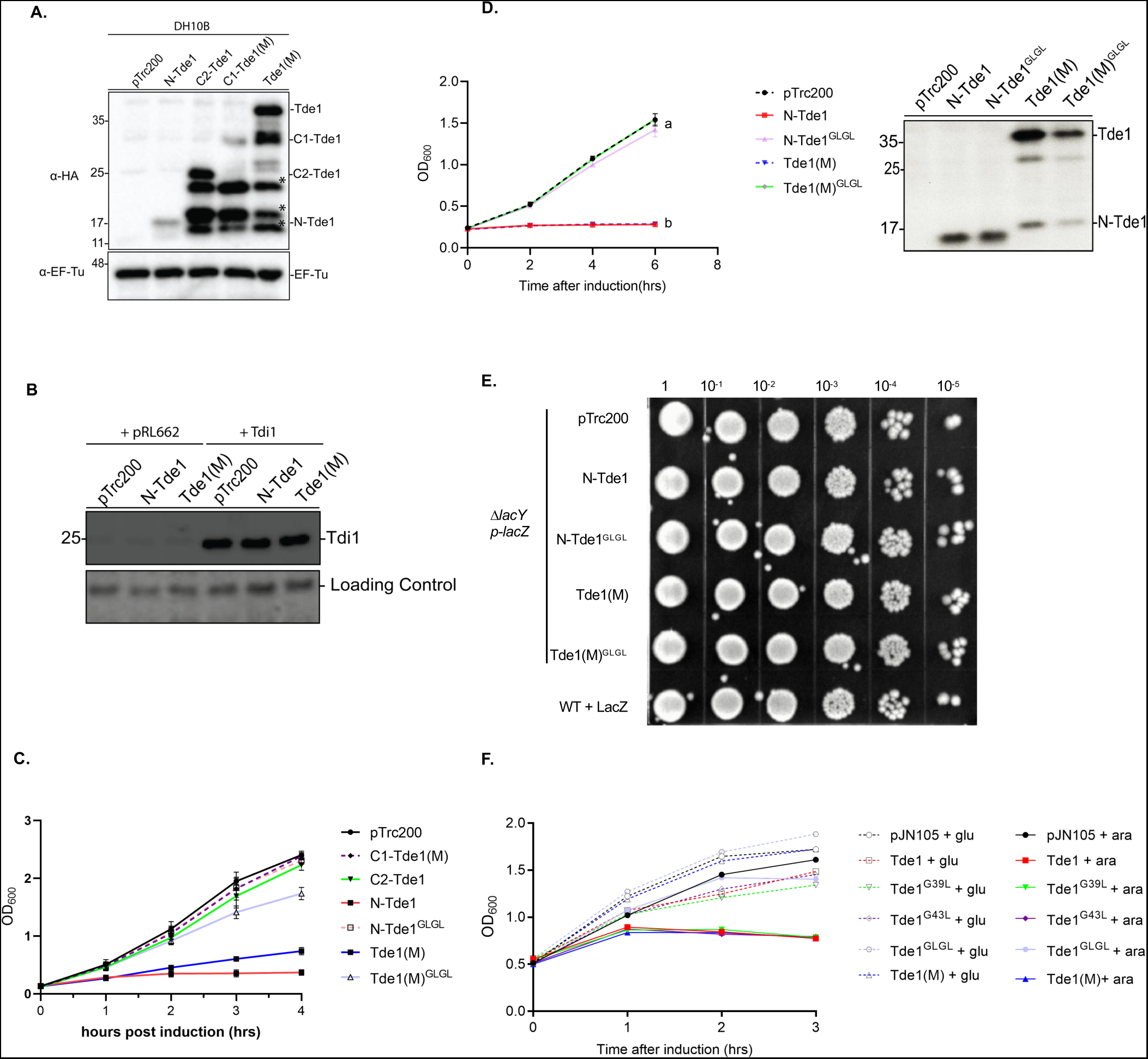
Growth inhibition assays in *A. tumefaciens* and *E. coli*. (A) Western blot for detection of the expression of HA-tagged Tde1 variants expressed in *E. coli.* *Other HA-tagged truncated bands, related to Figure 1B. (B) Western blot for the detection of Tdi1 from *E. coli* cells co-expressing HA tagged Tde1 variants and strep-tagged Tdi1, related to Figure 1C. (C) Growth inhibition assay of *E. coli* DH10B cells harbouring pTrc200 vector or each of its derivatives expressing Tde1 variants with IPTG inducible expression, monitored by OD_600_, related to Figure 2A. (D) Growth curve and western blot analyses of *A. tumefaciens* C58 Δ*tde1* carrying pTrc200 or its derivatives expressing HA-tagged Tde1 variants. The growth curve was detected every 2 hrs in 523 medium supplemented with 1 mM IPTG. Graphs show mean ± SD of three independent experiments. Different letters indicate statistically different groups of strains (p value <0.01). The proteins collected at end point (6 hr) were analysed for western blotting with antibody against HA. Representative results of three independent experiments were shown. Protein markers are indicated in kDa. (E) Viability assay for *E. coli* cells derived from the ONPG uptake assay after 1 hr IPTG induction, related to Fig. 2B. (F) The growth curve analysis of *E. coli* cells used for *in vivo* plasmid DNA degradation assay. The turbidity of *E. coli* BW25113 expressing Tde1 and its variants carried out for the *in vivo* plasmid DNA degradation assay were measured. The *E. coli* cells were supplemented with 0.5% glucose (glu) or 0.2% L-arabinose (ara) for the repression or induction of Tde1 and its variants. The OD_600_ values were measured by DEN-600 photometer (Biosan, Latvia) every hr.

**Figure S2.**
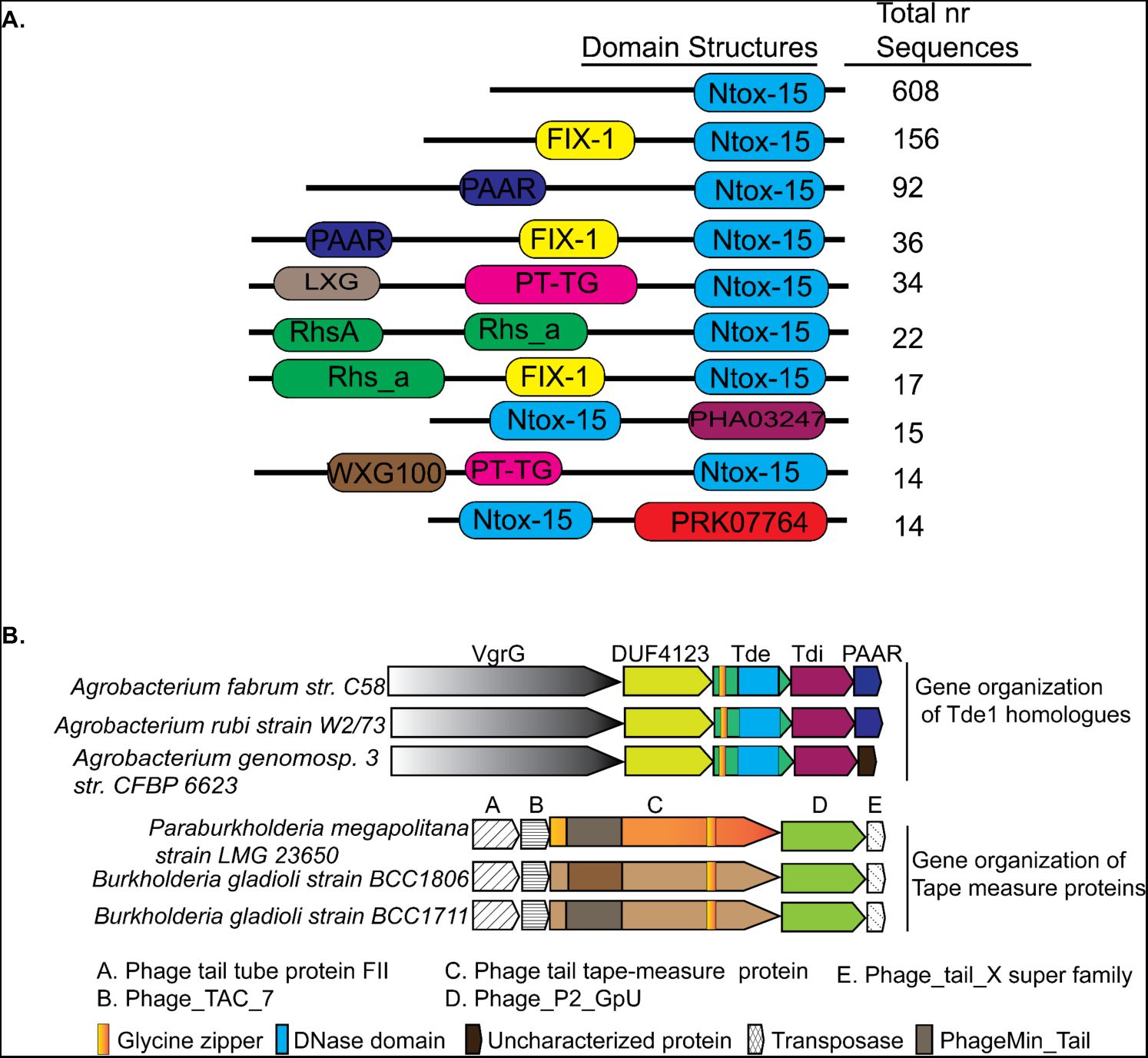
Domain architecture and genetic organizations of Tde homologues. (A) Domain organization of the Ntox15-containing proteins. Top 10 classes of the Ntox15-containing proteins are shown with the identifiable domains (not to scale). The number of proteins in each class were indicated on the left based on the information on June 29, 2022. The *A. tumefaciens* Tde1 belonged to the first class where the N-terminal region lacks an identifiable domain. (B) Genetic organizations of genes encoding Tde1 orthologues and Tape Measure Proteins (TMPs) with sequence similarity to N-terminus of Tde1. The proteins encoded from the upstream and downstream of *tde1* and *tmp* genes are shown with their identified domain organizations.

**Figure S3.**
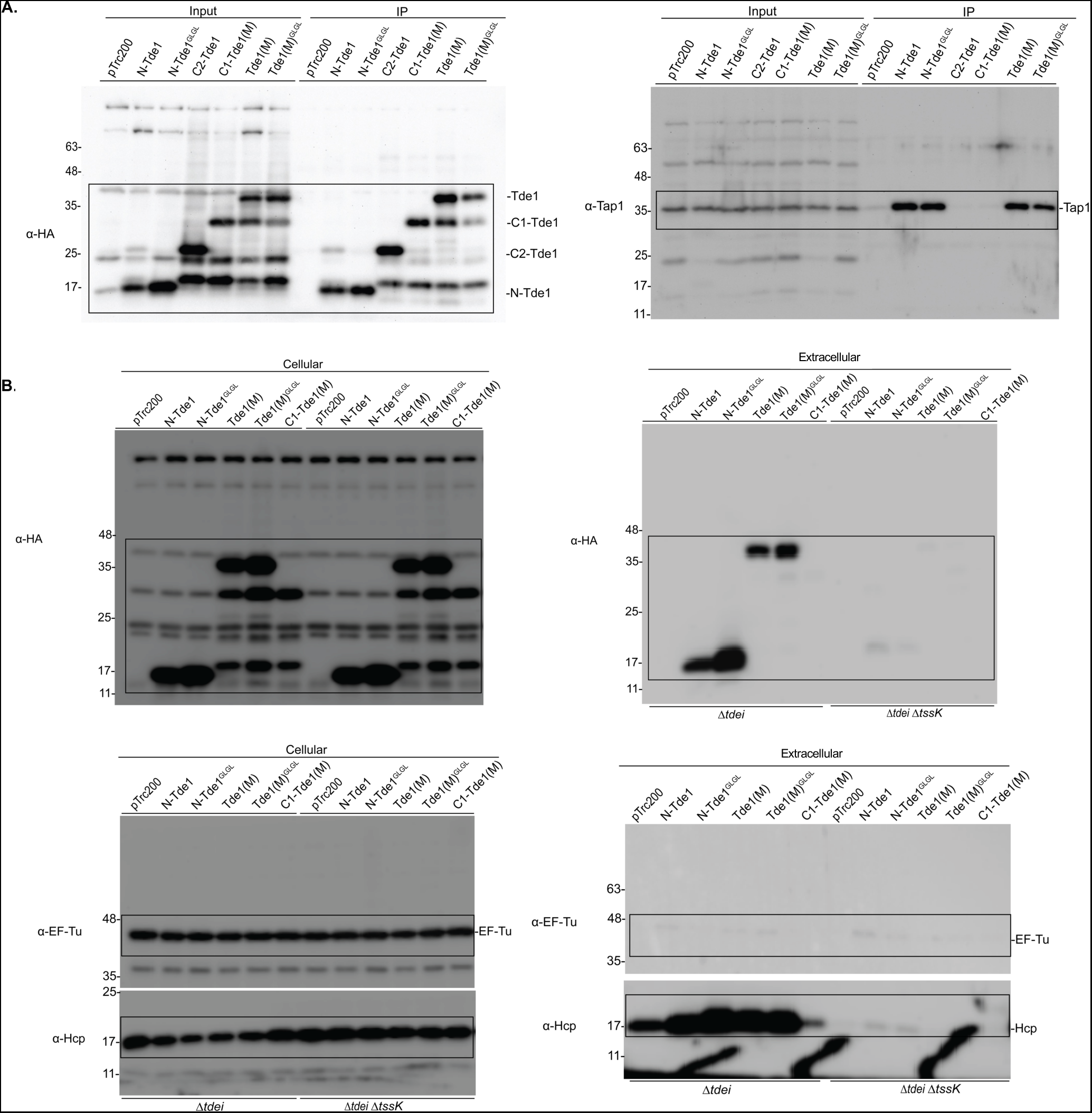
Full western blots of Co-IP and secretion assays. (A) Co-IP in *A. tumefaciens. A. tumefaciens* C58 Δ*tde1* harbouring pTrc200 vector or its derivatives expressing HA-tagged Tde1 variants. Anti-HA resin was used to coprecipitate the Tde1 variants and Tap1. (B) Secretion assay for HA-tagged Tde1 variants. Western blot for the cellular and extracellular fractions of *A. tumefaciens* C58 Δ*tdei* and Δ*tdei1ΔtssK* expressing the HA-tagged Tde1 variants. Hcp secretion was detected as a positive control for active T6SS secretion. Representative western blot results of three independent experiments were shown with antibody against HA, Hcp, or EF-Tu serving as a loading and non-secreted protein control. Protein markers are indicated in kDa. Box indicates the cropped area shown in Fig. 3.

**Fig. S4.**
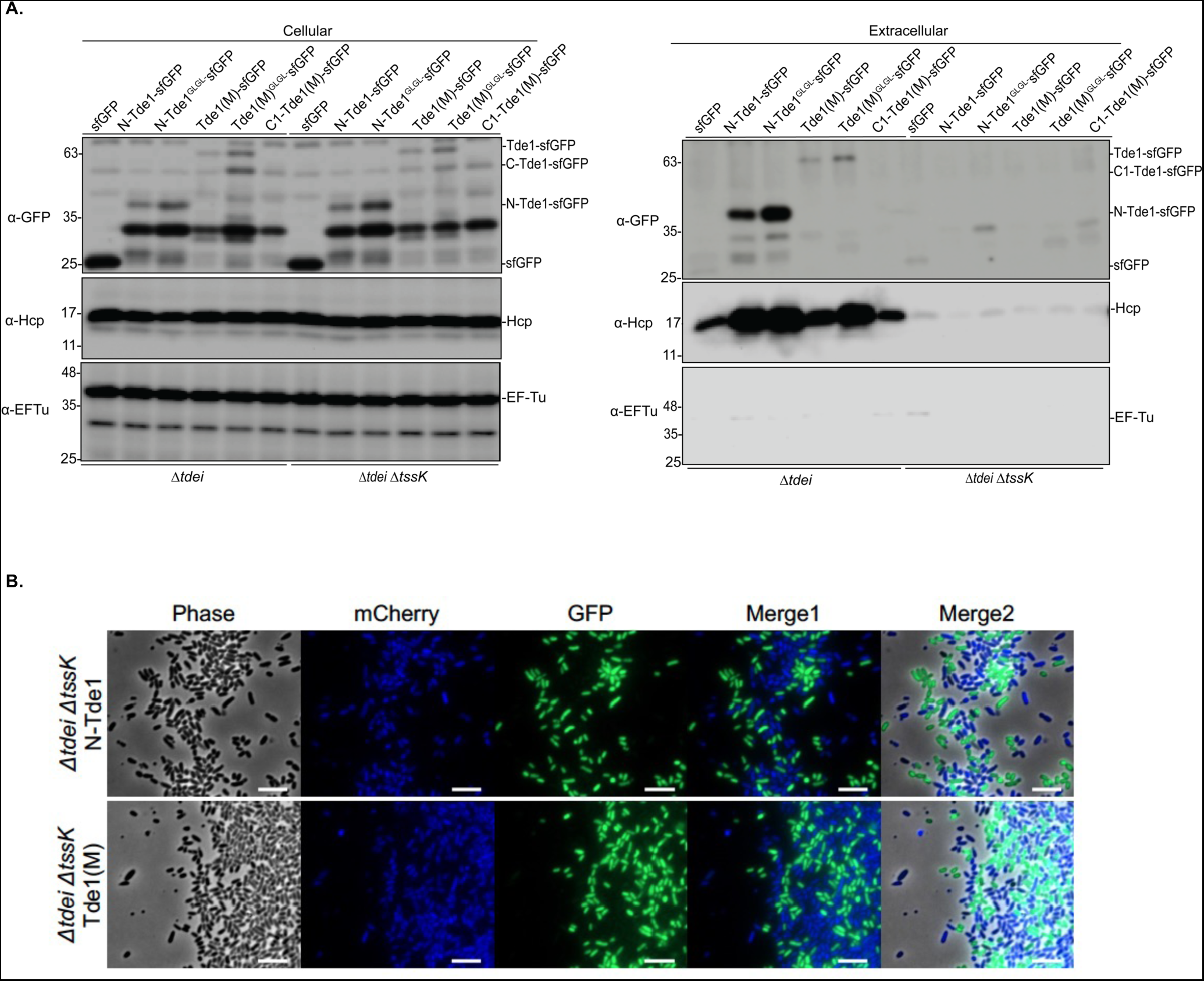
Fluorescence microscopy for negative controls of translocation assay. (A) Secretion assay for Tde1 variants fused with sfGFP. Western blot for the cellular and extracellular fractions of *A. tumefaciens* C58 Δ*tdei* and Δ*tdei1ΔtssK* expressing the Tde1 variants fused with sfGFP were detected by anti-GFP antibody. Representative western blot results of three independent experiments were shown with antibody against GFP, Hcp, or EF-Tu serving as a loading and non secreted protein control. Hcp secretion served as a positive control for active T6SS secretion. Protein markers are indicated in kDa. (B) *A. tumefaciens* C58 Δ*tdeiΔtssK* expressing N-Tde1-sfGFP or Tde1(M)-sfGFP (in green) and *E. coli* DH10B carrying mCherry (false coloured in blue) were co-cultured for 20 hrs. No cyan fluorescence with merged blue and green signals could be detected when attacker cells are T6SS-inactive, which served as negative controls for the translocation assay (Scale bar = 5μm).

**Figure S5.**
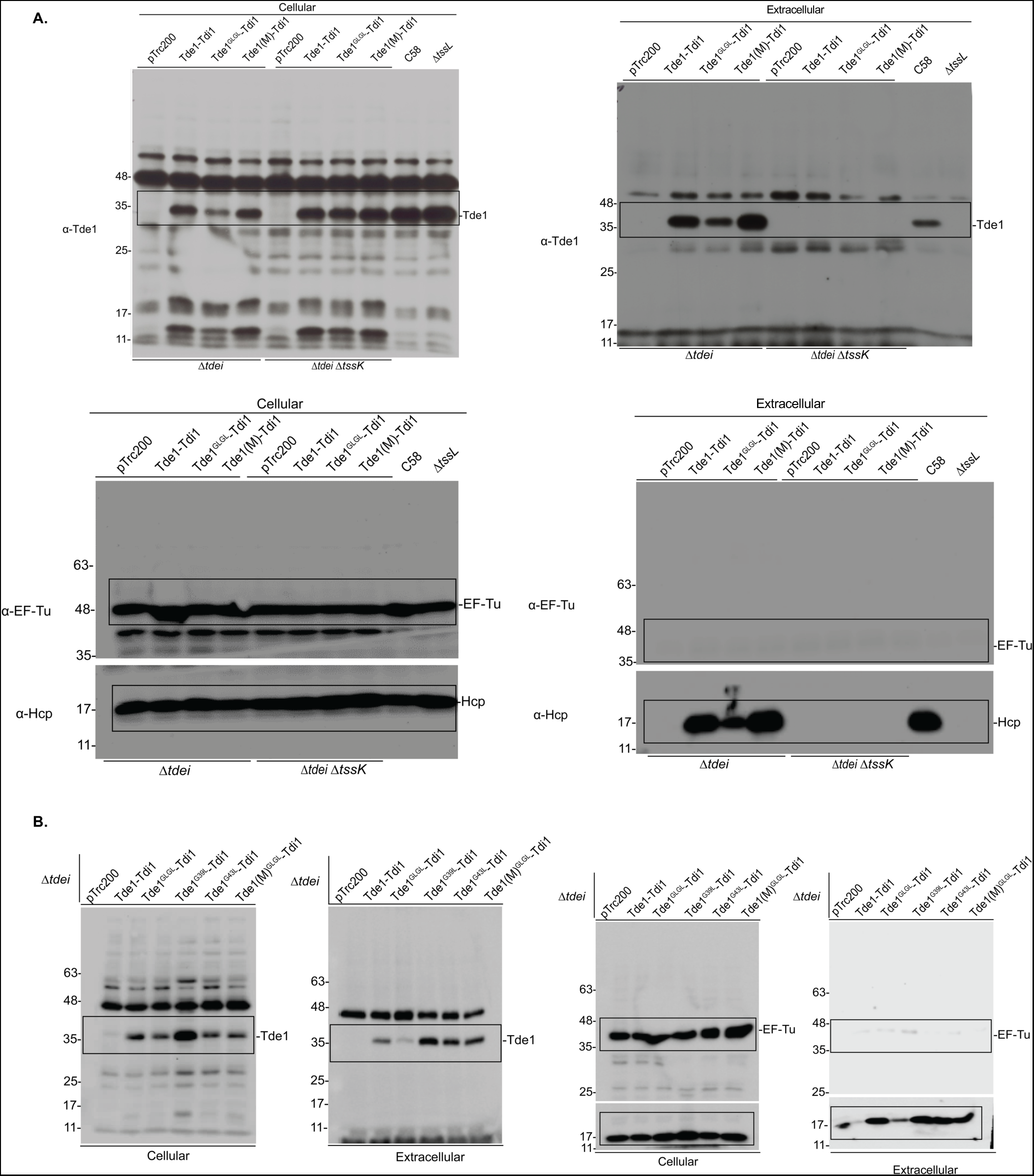
Full western blots of secretion assays. Secretion assay for Tde1 and its variants co-expressed with its immunity protein Tdi1 in *A. tumefaciens* C58 Δ*tdei* and Δ*tdeiΔtssK*. Western blots were detected with specific antibody against Tde1, Hcp, or EF-Tu serving as a loading and non-secreted protein control. Protein markers are indicated in kDa. Box indicates the cropped area shown in Fig. 5B, 5E.

**Figure S6.**
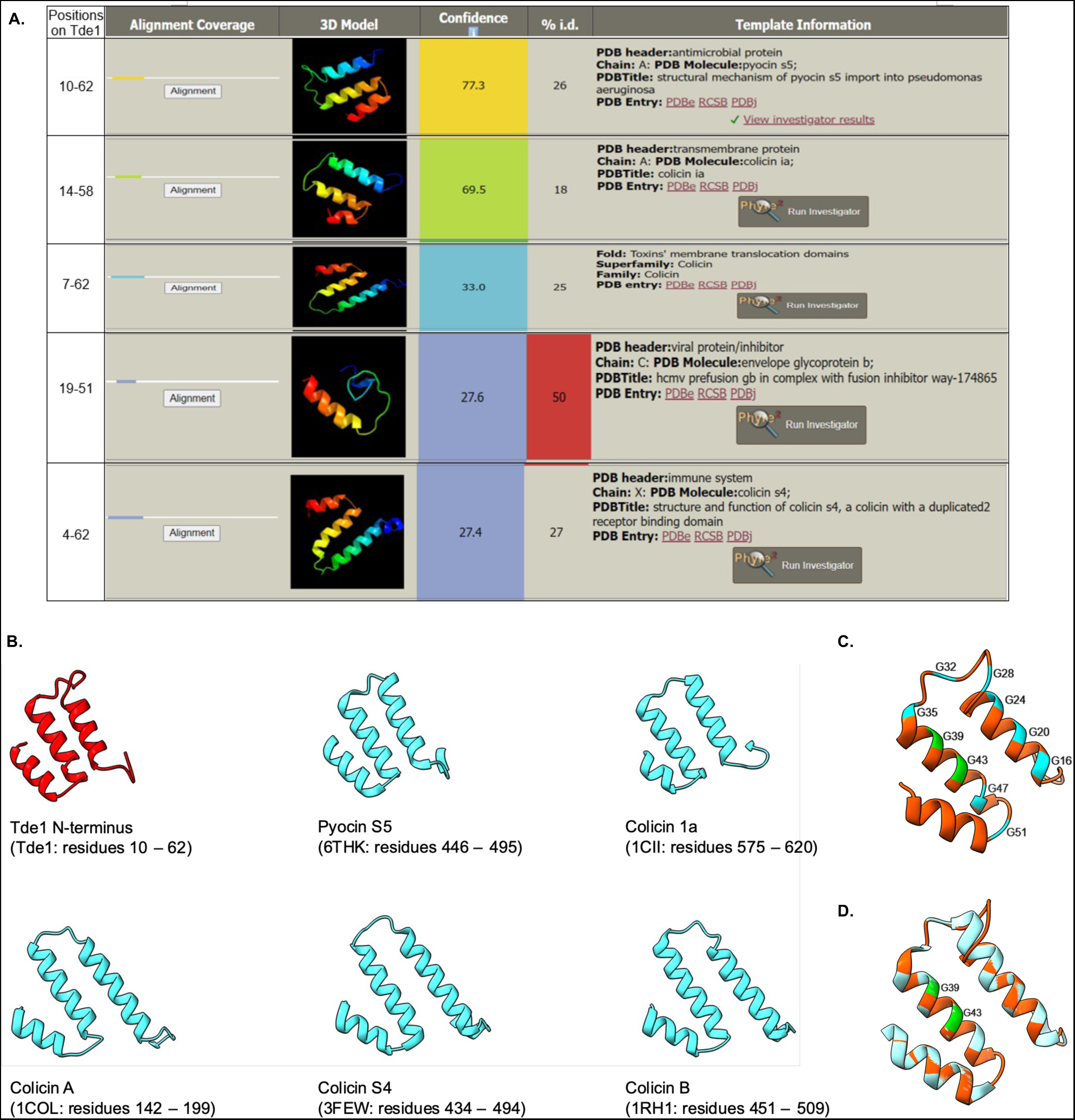
Structural prediction of the Tde1 N-terminus with similarity to pyocin S5 and colicins. (A) Predicted results of N-terminal Tde1 (1-97) as a query. (B) N-terminal Tde1 with structural similarity to pore-forming domain of the pyocin S5, colicines and other membrane perturbing proteins based on Phyre2 prediction. (C) Cartoon model of the Tde1 (residue 10 – 62) by using on the basis of the crystal structure of Pyocin S5 (PDB 6THK) with 77.3% of identity. Glycine residues of Tde1 were indicated. (D) Superimposition of N-Tde1 and pore-forming domain of pyocin S5. Tde1 N-terminus is in red and partially pore-forming domain of pyocin S5 is in teal, G^39^ and G^43^ in the putative glycine zipper motif are highlighted in green. All data were analyzed by Phyre2 server.

**Table S1.**
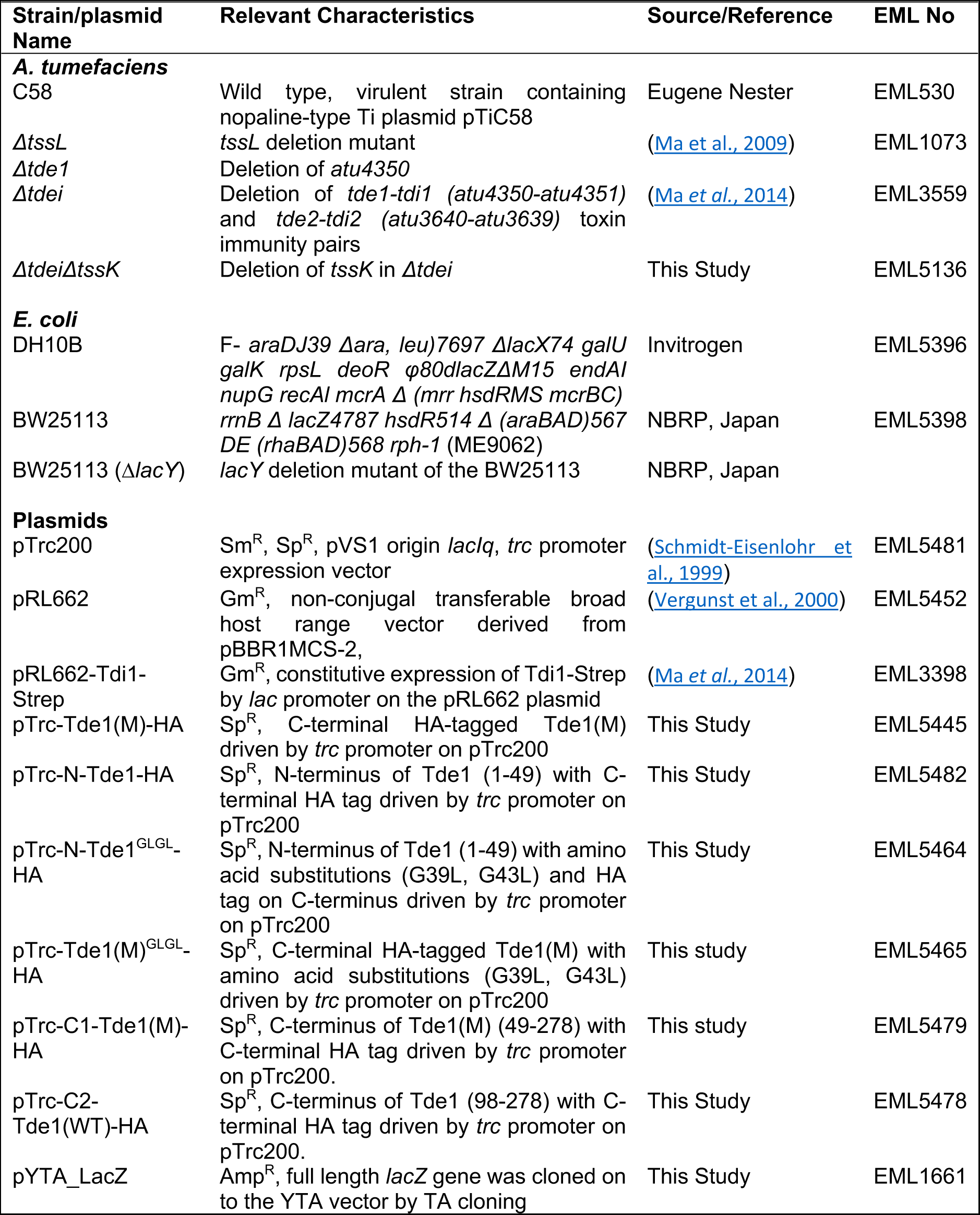
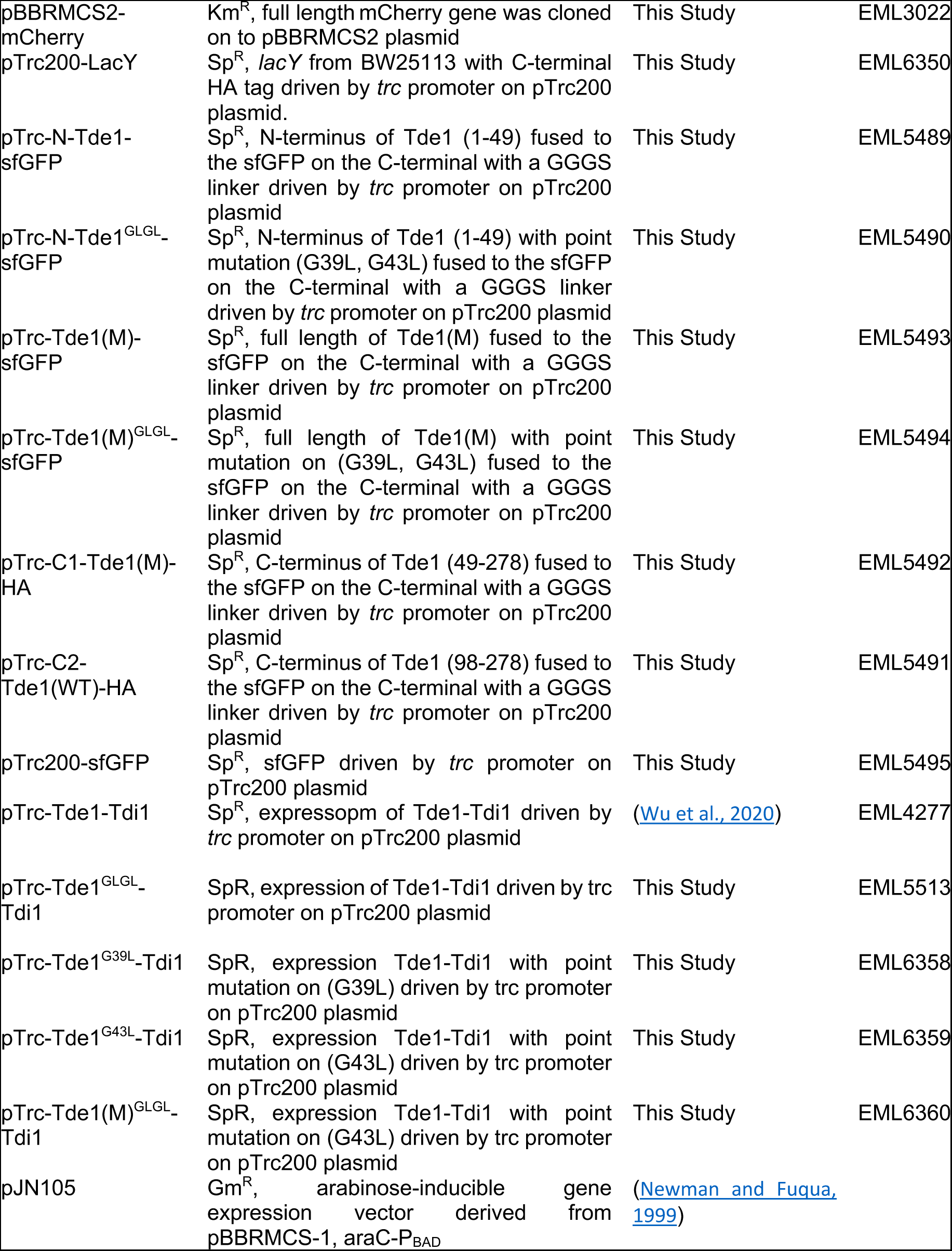
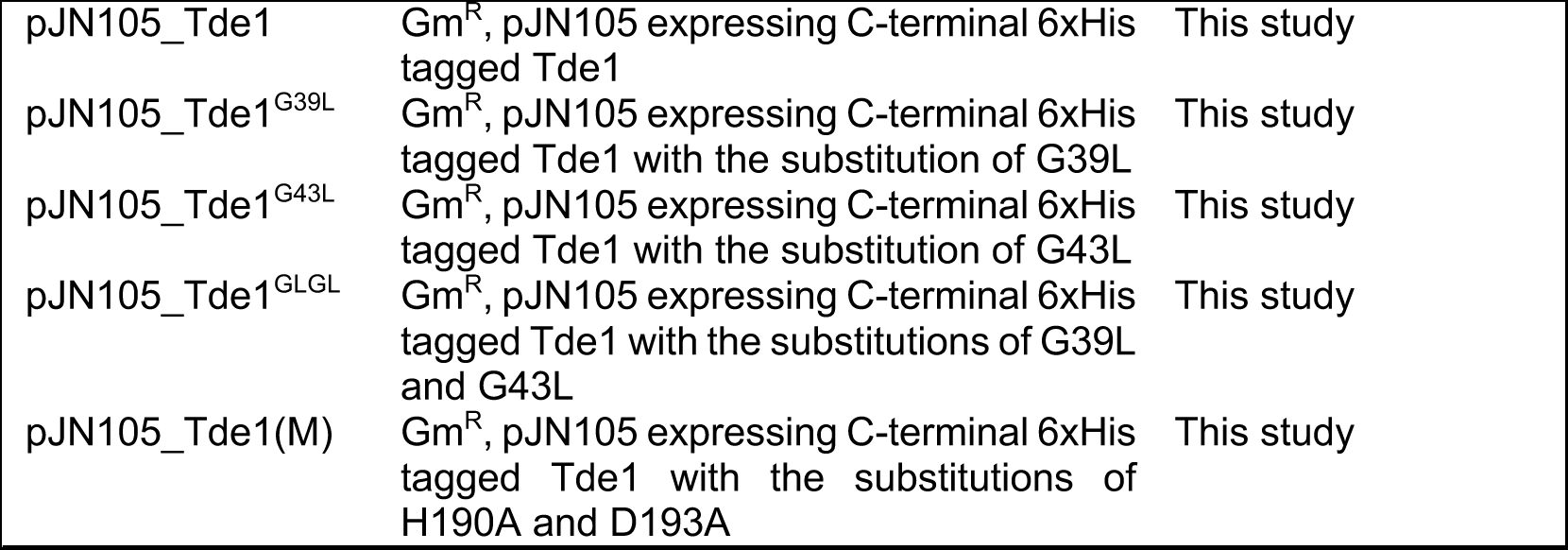
List of Strains and plasmids.

**Table S2.**
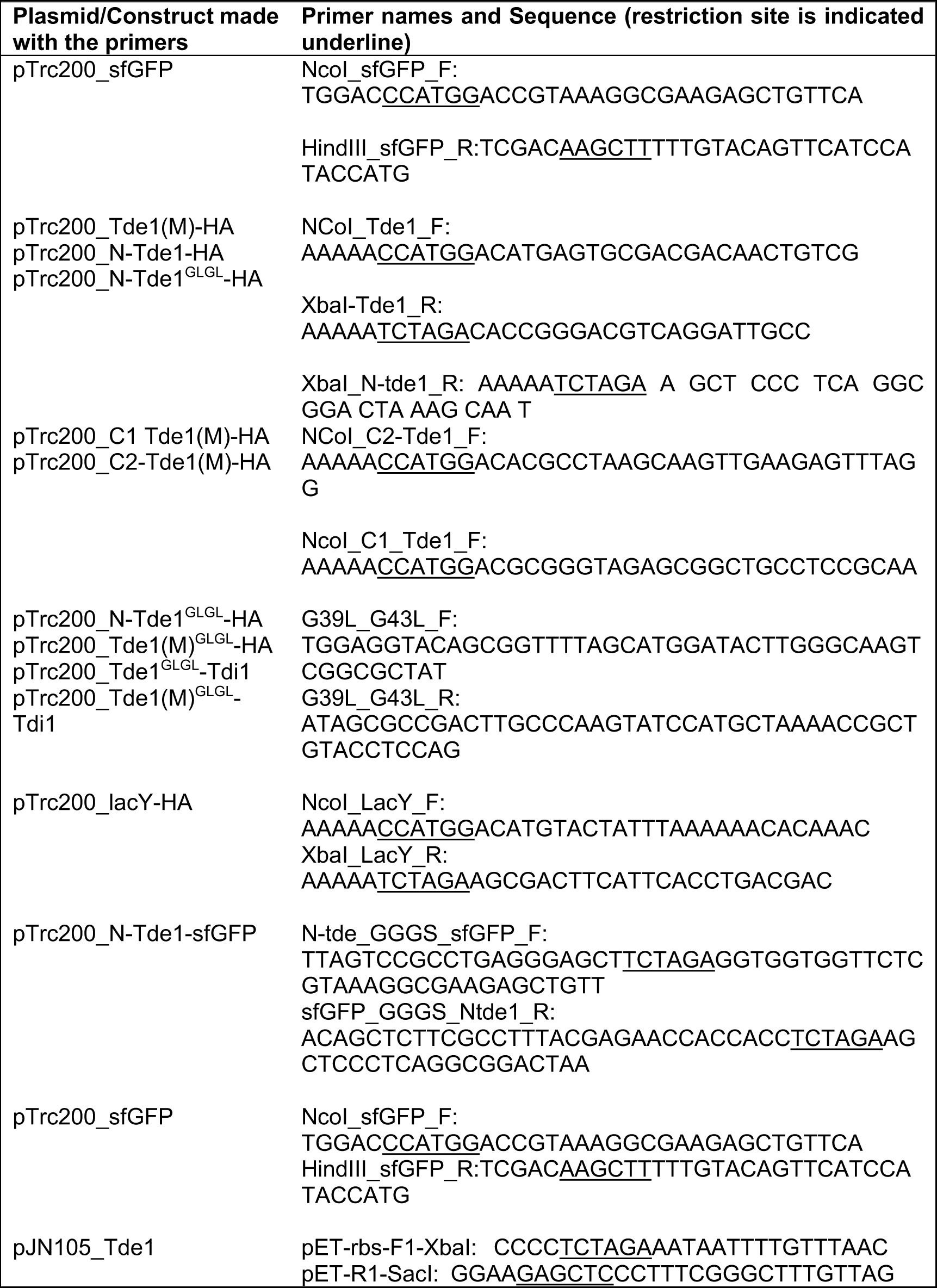
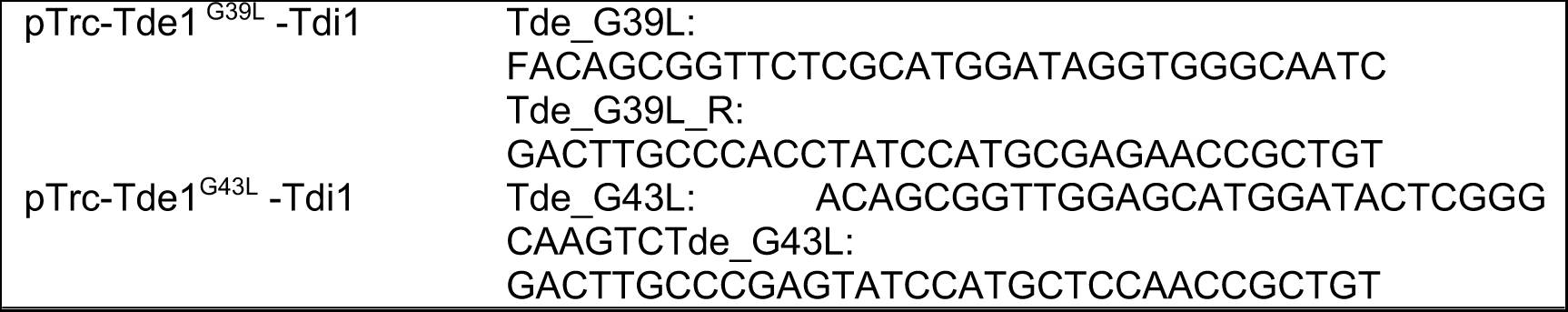
List of primers used to make constructs.

